# Precursor miRNAs are trafficked along axons associated with vesicles and locally processed to regulate growth cone steering

**DOI:** 10.1101/470393

**Authors:** Eloina Corradi, Antoneta Gavoci, Stephanie Strohbuecker, Michela Rocuzzo, Irene Dalla Costa, Archana Iyer, Simone Bridi, Gabriela Santoz Rodriguez, Cei Abreu-Goodger, Marie-Laure Baudet

## Abstract

Various species of non-coding RNAs (ncRNAs) are enriched in subcellular compartments but the mechanisms orchestrating their delocalization and their local functions remain largely unknown. We investigated both aspects using the elongating retinal ganglion cell axon and its tip, the growth cone, as models. We reveal that specific endogenous precursor microRNAs (pre-miRNAs) are actively trafficked, anchored to CD63-positive vesicles, to distal axons along microtubules. Upon exposure to the chemotropic cue Sema3A, pre-miRNAs are processed specifically within axons into newly synthesized mature miRNAs, which, in turn, silence the basal translation of TUBB3 but not of APP. At the organismal level, these mature miRNAs are required for growth cone steering and a fully functional visual system. Overall, our results uncover a novel mode of ncRNA transport from one cytosolic compartment to another within polarized cells. They also reveal that newly synthesized miRNAs are critical components of a ncRNA-based signaling pathway that transduces environmental signals into the structural remodelling of subcellular compartments.

**Highlights:**

- Precursor miRNAs are actively transported along axons to the growth cone tethered to CD63-positive vesicles
- Sema3A but not Slit2 induces the local biogenesis of specific miRNAs within axons
- Mature miRNAs are important for growth cone responsiveness *ex vivo* and the establishment of functional connections *in vivo*
- Newly synthesized miRNAs inhibit the basal translation of TUBB3 but not APP upon Sema3A exposure

## Introduction

Most cells are polarized, with an intracellular milieu partitioned into various organelles, cytosolic and membrane microdomains that accomplish specialized biological and regulatory functions. Neurons are highly polarized cells and their axons constitute a unique cellular outpost with distinctive autonomous function. During development, axons grow and generate a complex network of interconnected neurons. To establish these connections, the tip of the growing axon, the growth cone, is guided by attractant and repellent chemotropic cues *en route* to its target with exquisite precision. Axons must sometimes navigate a significant distance before reaching their final destination. In extreme examples such as large mammals, axons can even reach targets located meters away. The distance between growth cone and cell body poses a particular challenge to developing neurons because growth cones must be able to rapidly and accurately transduce environmental information to ensure highly precise directional steering, without the immediate intervention from the soma. To overcome this challenge, growth cones store but also locally produce and fine tune the levels of their own proteins through local protein synthesis (LPS), from a rich repertoire of mRNAs that is selectively trafficked there (Campbell and Holt, 2001; Davis et al., 1992; Wong et al., 2017; Zivraj et al., 2010).

Overall, mRNA localization and its corollary, local translation, are key mechanisms to create and sustain polarity by conferring functional autonomy to a variety of subcellular compartments including axons (Chin and Lécuyer, 2017). Recent evidence suggests that not only mRNAs but also various additional RNA species such as the small non-coding RNAs (ncRNAs) miRNAs (Kye et al., 2007; Lugli et al., 2008; Natera-Naranjo et al., 2010), linear long ncRNAs (Cabili et al., 2015), and circular RNAs (You et al., 2015) are delocalized to and enriched within subcellular outposts. The mechanisms of ncRNA transport to these compartments and the biological functions of local ncRNAs have, however, remained elusive. Gaining such fundamental insight using axons as a model is crucial because derailed axonal transport is a common factor in several neurodegenerative disorders for which there is currently no cure (Liu et al., 2012).

Dicer is a key component of the miRNA-induced silencing complex (miRISC) which is essential for the production of ~22 nt active, mature miRNAs from hairpin precursor miRNAs (pre-miRNAs) (Grishok et al., 2001; Hutvagner et al., 2001). Remarkably, numerous reports have revealed the presence of Dicer within growth cones (Aschrafi et al., 2008; Gershoni-Emek et al., 2018; Hancock et al., 2014; Hengst et al., 2006; Kim et al., 2015b; Vargas et al., 2016; Zhang et al., 2013). These findings raise the intriguing possibility that inactive pre-miRNAs are trafficked along axons to the growth cone and locally processed for function. Here, we first explore axonal pre-miRNA transport dynamics using a novel molecular beacon-based approach. We unravel that pre-miRNAs exploit a vesicle-mediated transport mechanism to reach the tip of the axon where they are subsequently stored. We then investigate the local function of pre-miRNAs within growth cones. We uncover that upon repellent cue exposure, specific pre-miRNAs are locally processed into active miRNAs that rapidly inhibit the basal translation of a selected transcript. This, in turn, induces directional steering *ex vivo* and *in vivo*, and promotes the development of a fully functional neuronal circuit. Collectively, these results reveal that miRNAs are trafficked tethered to vesicles to distal axons in an inactive form. At the growth cone, they are activated, on demand, to acutely inhibit basal translation of specific transcripts ultimately promoting growth cone steering and the highly accurate assembly of neuronal circuits.

## Results

### Dicer and pre-miRNAs are present within axons

Numerous studies have reported the presence of Dicer within mammalian growth cones by immunofluorescence in culture (Aschrafi et al., 2008; Gershoni-Emek et al., 2018; Hancock et al., 2014; Hengst et al., 2006; Kim et al., 2015b; Vargas et al., 2016; Zhang et al., 2013), but whether Dicer is also detected in the growth cones of lower vertebrates such as *Xenopus laevis* is so far unknown. We thus examined the distribution of Dicer in RGC axons in this species (Fig. 1A,B). We also examined the presence of Ago2, a key miRISC component required for miRNA-mediated silencing. A clear punctate Dicer- and Ago2-immunoreactive signal was observed in growth cones (Fig. 1B). Dicer is thus widely distributed in growth cones from many different cells types and species suggesting an important and universal local role in this compartment.

**Figure 1:**
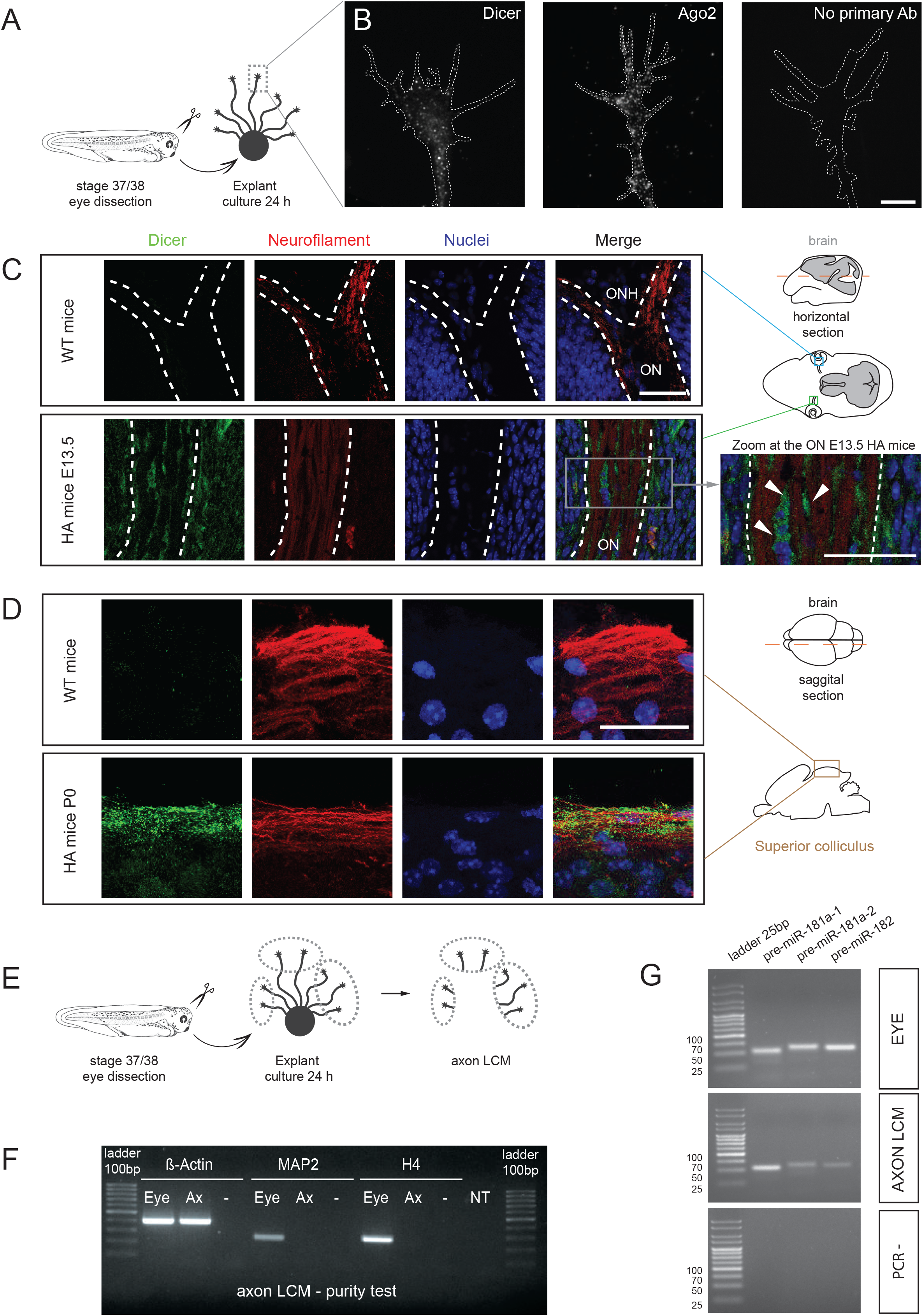
Dicer, Ago2 and pre-miRNAs are localized in RGC axonal compartments. **(A, E)** Schematic representation of the experimental protocol. **(B)** Representative *Xenopus* growth cones stained with anti-Dicer and anti-Ago2 antibodies. Negative control: no primary antibody (Ab). **(C)** Representative E13.5 mice brain cross-section (dashed red line in the schematic) comprising the optic nerve (ON) stained with anti-neurofilament and anti-HA antibodies to detect RGC axons and (FLAG-HA_2_)-Dicer, respectively. Note the absence of HA signal in wild type (WT) mice. The white dashed delineates the ON. Zoom of the triple stained ON (right panel): Dicer signal is detected inside ON cells surrounding axon bundles but not in axons *per se* (arrowheads). **(D)** Representative P0 superior colliculus (sagittal sectioning of P0 brains along the dashed red line in the schematic). Note the presence of Dicer within neurofilament-marked RGC axons. **(F)** RT-PCR performed on RNA extracted from axons collected by LCM (Ax) or from eyes. β-Actin mRNA is present both in eye and axons, while MAP2 and H4 are present only in the eye sample, suggesting the absence of contamination from cell bodies or other cells in LCM axonal samples. **(G)** RT-PCR performed on RNA extracted from eyes or from LCM isolated axons. Statistics: (**C, D**) n=3 independent experiments. Abbreviations: WT, wild type; HA, (FLAG-HA_2_)-Dicer; E13.5, embryonic day 13.5; P0, post-natal day 0; ONH, optic nerve head; ON, optic nerve; LCM, laser capture microdissection; Ax, axonal sample; Eye, stage 37/38 eye; -, no template control of the PCR; NT, no template RT control. Scale bars: 10 µm (B); 30 µm (C, D). *See also Figure S1*.

To confirm the *bona fide* expression of Dicer within axons without the reliance on anti-Dicer antibody, we used an endogenous epitope tagged (FLAG-HA_2_)-Dicer knock-in mouse allele (Much et al., 2016) (Fig. S1A,B) and investigated the distribution of Dicer in RGC axons by immunofluorescence with anti-HA antibody *in vivo*. Specificity of the anti-HA antibody was confirmed by the absence of signal in WT mice (Fig. 1C,D; Fig. S1C) and as expected, Dicer was detected within the cytoplasm of retinal cells (Much et al., 2016) (Fig. S1C). At E13.5, when RGC axons are reaching the chiasm (Bovolenta and Mason, 1987), Dicer was not detected in axons marked with anti-neurofilament antibody but appeared in cells within the optic nerve head (Fig. 1C, arrowheads). By P0, Dicer was detected within RGC axons where HA- and axonal neurofilament-associated signals clearly overlapped (Fig. 1D). At this stage, axons start innervating their target centers, the superior colliculus and lateral geniculate nucleus (Godement et al., 1984). This strongly suggests that Dicer, and by extension newly synthesized miRNAs, may act within RGC axons not during the period of axon pathfinding but during the process of axon targeting. Overall, these results indicate the presence of Dicer within mammalian axons *in vivo* during a specific developmental window.

The detection of Dicer at the growth cone suggests that miRNAs could be locally produced within this compartment. If this were the case, inactive hairpin miRNA precursors should be detected in axons and growth cones. Our previously published dataset of small RNAs from axons and growth cones (Bellon et al., 2017) shows the presence of sequences corresponding to the loop region of 42 pre-miRNAs (Fig. S1D). Pre-miRNA-182, pre-miR-181a-1 and pre-miR-181a-2 were amongst the 9 most abundant pre-miRNAs in axons, as determined by the number of mapped reads (Fig. S1D). To validate the presence of these precursors within axons, we performed RT-PCR from axonal RNA. The axonal RNA was collected via laser capture microdissection (LCM) from RGC axons derived from stage 37/38 whole eye explant cultures (Fig. 1E). This approach yields pure axonal RNA as indicated by the presence of the well-described axonal marker *β*-Actin and the absence of both the dendritic marker MAP2 and the nuclear marker histone H4 (Bellon et al., 2017) (Fig. 1F). PCR confirmed the presence of pre-miR-181a-1, pre-miR181a-2 and pre-miR-182 in eyes and axons (Fig. 1G). These two precursors showed different relative levels in whole eyes and isolated axons, with miR-181a-1 precursors being 5.93 significantly less abundant than miR-181a-2 precursors in whole eyes (Fig. S1E), yet 2.54 more abundant in isolated axons (Fig. S1F). This indicates that pre-miR-181a-1 might be preferentially targeted to axons and growth cones.

### Molecular beacons are new, specific tools to detect endogenous pre-miRNAs in cells

Pre-miRNAs may be located in axons due to passive diffusion. Alternatively, they may be actively and directionally transferred along axons to growth cones to exert a compartmentalized function. To address this, we examined the dynamic transport of pre-miR-181a-1 along axons. We focused on pre-miR-181a-1 because it is more abundant in axons than pre-miR-181a-2 (Fig. S1F).

In order to specifically examine the behavior of native pre-miRNAs within the cell, we developed a novel visualization approach to track endogenous pre-miRNAs using molecular beacons (MBs). MBs are single-stranded oligonucleotide probes which fluoresce only when hybridized to their target (Santangelo et al., 2006) (Fig. 2A). The MB backbone and sequence was carefully designed to maximize the probe’s 1) stability, 2) signal-to-noise ratio, and 3) specificity within the cell (see Methods for details). Tests *in vitro* indicated that the MB is stable in presence of DNAse I or total eye lysate (data not shown). Denaturation profiles confirmed the suitable thermodynamic characteristics of our MB design: low fluorescence at low temperature and a melting temperature (Tm) of 58°C, appropriate for our working temperature of 20°C (Simon et al., 2010; Søe et al., 2011) (Fig. 2C, black line). A thermal denaturation assay (Fig. 2B) revealed increased fluorescence levels at low temperatures in the presence of total RNA extracted from stage 40 whole eyes or *in vitro* transcribed pre-miR-181a-1, consistent with hybridization between the MB and synthetic or endogenous pre-miR-181a-1 (Fig. 2C; Fig. S2A). This confirms that our MB is able to specifically detect pre-miR-181a-1 *in vitro*.

**Figure 2:**
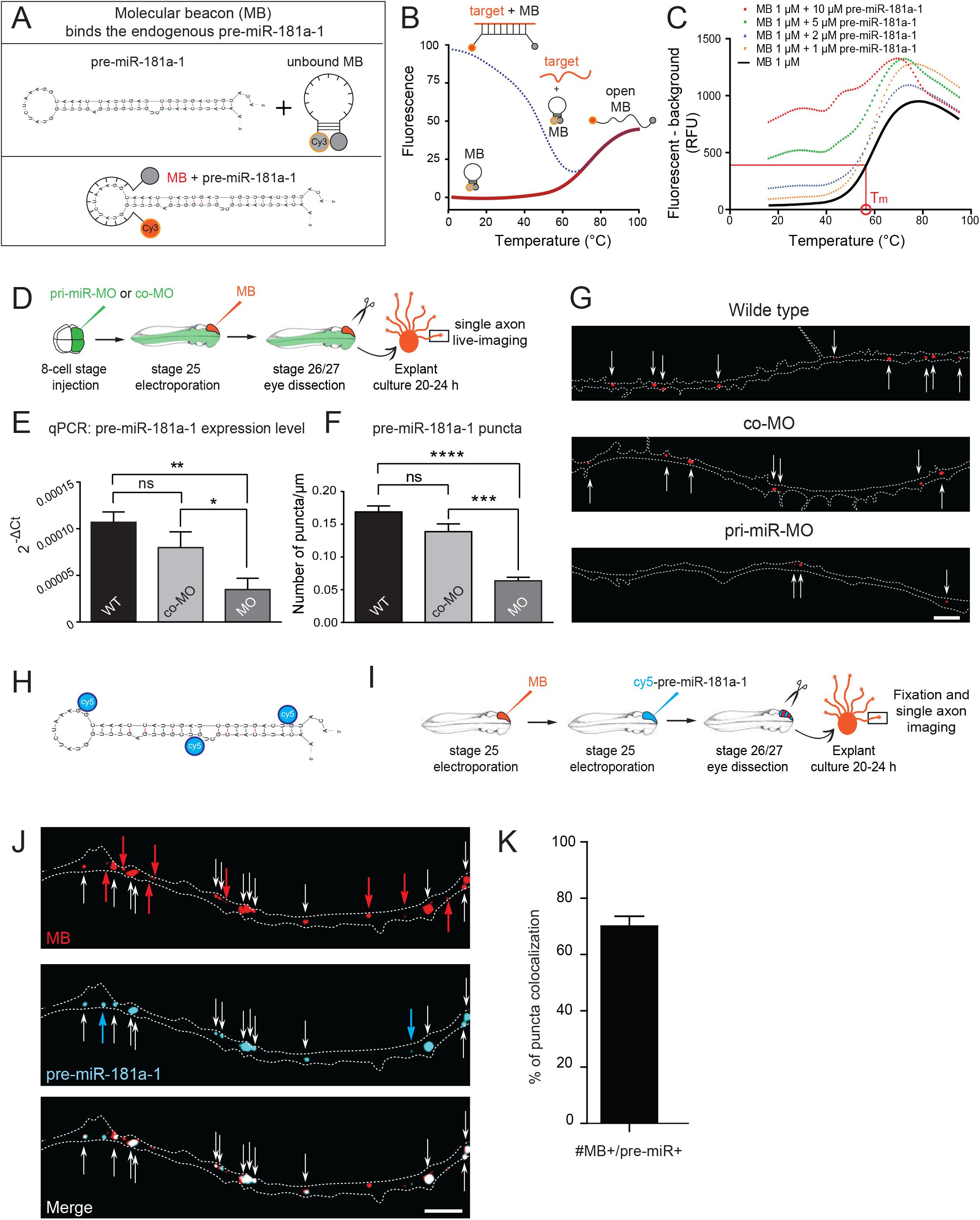
Pre-miR-181a-1 trafficked in RGC axons. **(A)** Schematic of molecular beacon (MB), pre-miR-181a-1 and their hybridization complex. **(B)** Schematic of thermal denaturation profile of the MB, in which the fluorescence signal is recorded as a function of temperature, both in absence (solid line) and presence (dashed line) of a target. **(C)** Thermal denaturation of 1 µM MB, in the absence (solid line) and presence (dashed line) of increasing concentration of pre-miR-181a-1 RNA target (1, 2, 5, 10 µM). MB melting temperature T_m_ is indicated in red in the plot. Note that the decrease fluorescence observed at the highest temperature is probably due to the fact that cy3 quantum yield (Φ) is sensitive to high temperature (Φ decreases by 70 % at 65°C compared to RT). Each melting curve represents the average of three separate experiments. **(D)** Schematic representation of the experimental protocol. Concentrations used: MB (5 µM), co-MO (250 µM), pri-miR-MO (250 µM), 226 ng/µl cy5-pre-miR-181a-1. **(E)** Quantification of the expression levels of pre-mir-181a-1 using the 2^^(-ΔCt)^ method and U6 as normalizer from small total RNA fraction (< ~150 nt). **(F)** Total number of MB puncta normalized to axon length (µm). **(G)** Representative axons. MB puncta are indicated (white arrow). Axons is delineated by the dashed lines. **(H)** Schematic representation of cy5-labeled pre-miR-181a-1. **(I)** Schematic representation of the experimental protocol. **(J)** Representative image of RGC axons derived from MB and cy5-pre-miR-181a-1 co-electroporated retina. White, red and blue arrows indicate, respectively, co-localized, single MB, and single pre-miR-181a-1 puncta. **(K)** Frequency (in percentage) of puncta co-localization between MB and cy5-pre-miR-181a-1 (#MB+/pre-miR+). Values are mean ± SEM (E, F, K). Statistics: * p<0.05, ** p<0.01, *** p<0.001 and **** p<0.0001. **(E)** Data were normally distributed (Shapiro-Wilk test), one-way ANOVA followed by Tukey’s multiple comparison post-hoc test, n=3 independent experiments. **(F)** Data were not normally distributed (Shapiro-Wilk test), Kruskal Wallis followed by Dunn’s multiple comparison post-hoc test. Total numbers of puncta analyzed: 928 (WT); 226 (MO); 208 (co-MO); from 61 axons (WT); 15 axons (co-MO); 35 axons (MO); n=4 independent experiments. (**K**) Total numbers of puncta analyzed: 354 (MB); 337 (cy5-pre-miR-181a-1); 32 single axons; n=5 independent experiments. Abbreviations: MB, molecular beacon; co-MO, control morpholino; pri-miR-MO, morpholino blocking pre-miR-181a-1 processing by targeting the Drosha cleavage site; ns, not significant. Scale bars: 5 µm (G, J). *See also Figure S2*.

We further tested MB specificity *ex vivo* using two strategies. First, we examined whether MB is still detectable when endogenous pre-miR-181a-1 is knocked down. To this end, we microinjected to 8-cell stage embryos morpholino (MO) designed to block Drosha cleavage and therefore reduce the levels of pre-miR-181a-1 (Fig. 2D). We named this MO pri-miR-MO. Pri-miR-MO induced a significant 56.29 % average decrease of axonal pre-miR-181a-1 levels compared to control MO (co-MO), as measured by qPCR (Fig. 2E). Consistently, the number of MB puncta was significantly decreased by an average 58.45 % within axons in pri-miR-MO vs co-MO treated embryos (Fig. 2F,G). We did not observe significant differences between WT and co-MO (Fig. 2E-G). Second, we examined whether cy3-labeled MB co-localized with exogenous cy5-labeled pre-miR-181a-1 (Fig. 2H) serially electroporated into eye primordia (Fig. 2I). Electroporation of MB or cy3-pre-miR-181a-1 was not toxic to the eye or to axons or growth cones (Fig. S2B,C). We detected 70.14 ± 3.51 % MB puncta co-localized with cy5-labeled pre-miR-181a-1 puncta, and 74.86 ± 3.55 % pre-miR-181a-1 puncta co-localized with MB puncta within axons (Fig. 2J,K). This suggests that the MB can specifically recognize pre-miR-181a-1 *ex vivo*.

### Endogenous and exogenous pre-miRNAs are actively trafficked along axons

We then characterized the endogenous pre-miR-181a-1 trafficking dynamics in RGC axons by live imaging following targeted electroporation of MBs into retinal cells (Fig. 3A). MB-labeled pre-miR-181a-1 puncta were detected throughout the entire length of growing RGC axons and accumulated within the central domain of growth cones (Fig. 3B). This confirms axonal translocation of endogenous pre-miR-181a-1 to the growth cone. Time-lapse acquisitions were quantitatively analyzed by transforming the movies into kymographs (Fig. 3C). To characterize pre-miRNA transport behavior, puncta’s velocity was calculated by considering the average speed of its segmental components. Equal frequencies of anterogradely and retrogradely moving puncta were detected (Fig. 3D), suggesting the existence of a balanced bidirectional transport. We next analyzed stationary (<0.2 μm/s), moving (0.2-0.5 μm/s) and fast moving puncta (>0.5 μm/s) (Leung et al., 2018; Maday et al., 2014) separately. No difference was observed between anterograde and retrograde average velocity for puncta slower than 0.5 μm/s (0.2 μm/s: antero, 0.066 ± 0.007 μm/s; retro, 0.084 ± 0.008 μm/s; 0.2-0.5 μm/s: antero, 0.33 ± 0.02 μm/s; retro, 0.32 ± 0.01 μm/s). However, fast (> 0.5 μm/s) anterogradely moving puncta were significantly faster on average than their retrograde counterpart (antero: 1.79 ± 0.08 μm/s versus retro: 1.16 ± 0.06 μm/s) (Fig. 3E). This difference was also reflected in a bimodal velocity distribution above 0.5 μm/s with faster velocities reached by anterogradely moving puncta (up to 3.4 μm/s) compared to retrogradely moving puncta (up to 2.6 μm/s) (Fig. S3A). Within axons, kinesin and dynein molecular motors are known to actively move cargos anterogradely and retrogradely, respectively, with a velocity of at least 0.5 μm/s (Maday et al., 2014). Contrary to the highly processive unidirectional motility of kinesins, dynein makes frequent back and side steps (Maday et al., 2014). The slower pre-miRNA retrograde transport that we observe at speeds above 0.5 μm/s may thus reflects dynein’s unique property. No significant difference in puncta directionality (Fig. 3D) and velocities (Fig. 3E; Fig. S3C,D) between endogenous and exogenous cy3-labeled pre-miR-181a-1 (Fig. 3A) was observed (see also Movie 1 and Movie 2). This strengthens the notion that the MB specifically detects pre-miR-181a-1 in axons and that both MB and cy3-pre-miR-181a-1 can be used interchangeably. Overall, this indicates that pre-miRNA molecules are transported along axons from the soma to the growth cone but also from the growth cone back to the soma at a range of velocities consistent with molecular motor-mediated trafficking (Maday et al., 2014).

**Figure 3:**
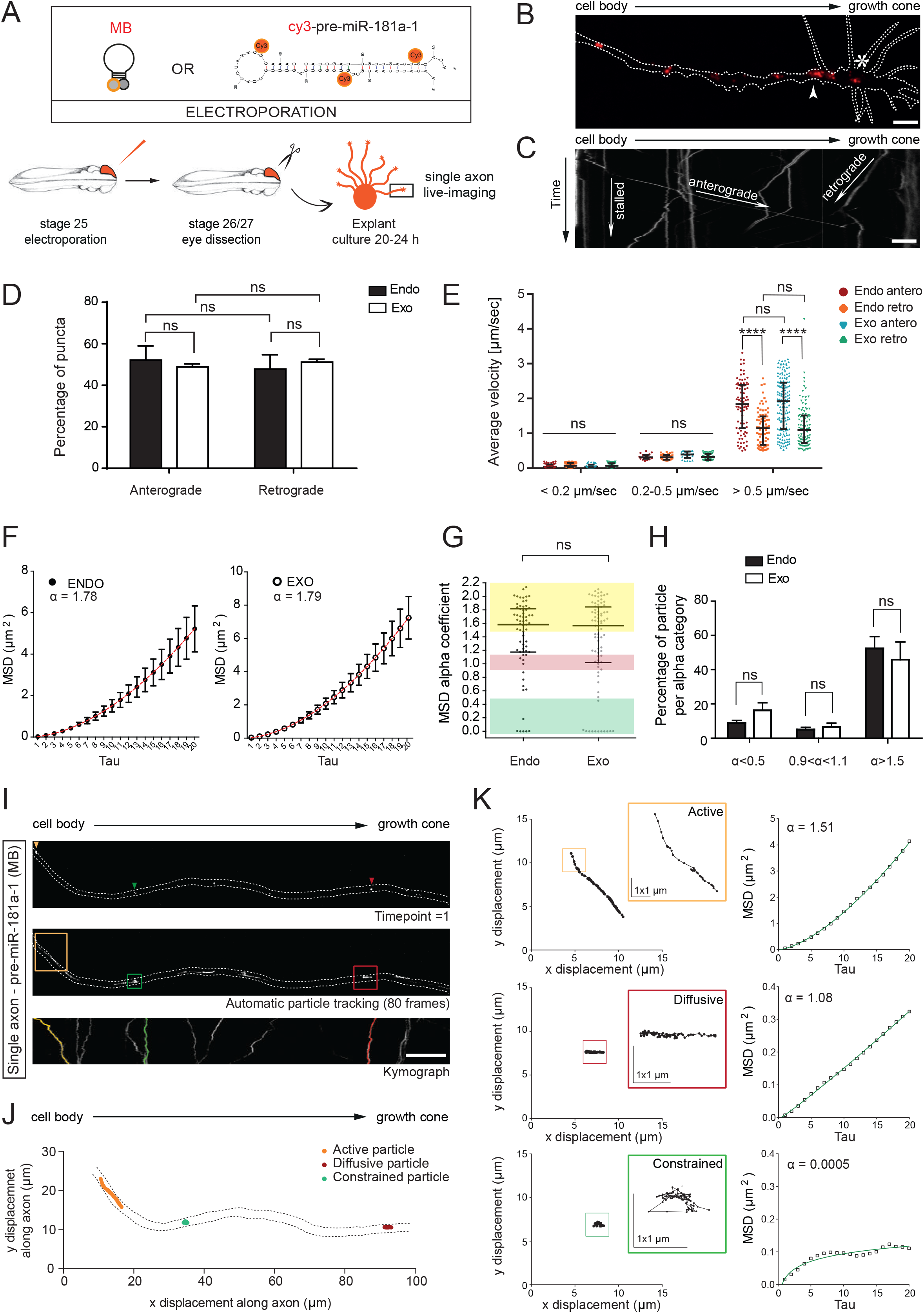
Pre-miR-181a-1 is actively trafficked in RGC axons. **(A)** Schematic representation of the experimental paradigm. MB was electroporated at 5 µM and cy3-pre-miR-181a-1 at 200 ng/ul. **(B)** Representative image of a single distal axon from MB-electroporated retina. Dashed white line delineates the axon tipped with a growth cone. **(C)** Illustrative kymograph generated from an acquired time-lapse. **(D)** Frequency distribution (in percentage) of anterogradely and retrogradely transported MB (endo) and cy3-pre-miR-181a-1 (exo) puncta along the RGC axon shaft. **(E)** Average velocity of MB (endo) and cy3-pre-miR-181a-1 (exo) puncta. Single dot corresponds to one puncta. **(F)** MSD data for MB (endo) and cy3-pre-miR-181a-1 (exo) tracked particles were fitted with an anomalous diffusion model and α thus calculated. The corresponding fitting is shown in red. **(G)** MSD alpha coefficient distribution for each single MB (endo) and cy3-pre-miR-181a-1 (exo) tracked particles. α>1.5, active particles in yellow; 0.9<α<1.1, diffusive particle in red; α<0.5, confined particles in green. **(H**) Relative frequency distribution (percentage) of confined (α<0.5), diffusive (0.9<α<1.1) and active (α>1.5) MB (endo) and cy3-pre-miR-181a-1 (exo) tracked particles. **(I, J)** Representative tracked active (yellow), diffusive (red), constrained (green) particles from one illustrative axon from MB electroporated retina. The dashed white (I) or black (J) lines delineate the axon. **(J)** The x-y trajectory over the 80 consecutive acquisition frames for each motion type is represented within the axon. Note the difference in trajectory length according to the particle type. **(K)** High spatial resolution of x,y displacement for each motion type and corresponding MSD plot with associated fitting and α value. Note the striking difference in trajectories according to particle type. Only a small portion of the actively moving particle trajectory is shown in the yellow inset (K). Values are mean ± SEM (D, F, H) or median with interquartile range (E,G). Statistics: **** p<0.0001. **(D-E)** Total number of analyzed particles: 353 (endo); 484 (exo). Endo n=3; exo n=4 independent experiments. **(D)** Two-way ANOVA followed by Sidak’s multiple comparisons post-hoc test. **(E)** Two-way ANOVA followed by Tukey’s multiple comparison post-hoc test. **(F-H)** Total number of analyzed particles: 67 (endo); 82 (exo). Endo n=3; exo n=4 independent experiments. **(G)** Data are not normally distributed (Shapiro-Wilk test), two-tailed Mann Whitney test. **(H)** Two-way ANOVA followed by Tukey’s multiple comparison post-hoc test. Abbreviations: ns, not significant; MB, molecular beacon; Endo, endogenous; Exo, exogenous. Scale bars: 5 µm (B, C); 10 µm (I); 1 µm x 1 µm (K). *See also Figure S3*.

To address whether moving (≥ 0.2 μm/s) endogenous pre-miRNAs were driven by active transport or passive diffusion, we performed mean square displacement (MSD) analysis (see Methods for details). The MSD data were fitted with an anomalous diffusion model: MSD = Aτ ^α^ + B (Eq1) (Otero et al., 2014). Trajectories were conservatively classified as actively driven (α>1.5), diffusive (0.9<α<1.1) or confined (α<0.5) depending on the value of the coefficient obtained from the fitting of Eq1 to the MSD data (Otero et al., 2014). We obtained α = 1.78 (endogenous) for the combined trajectories of all particles (Fig. 3F), suggesting that moving pre-miRNAs were overall actively trafficked along axons. We analyzed the α distribution of individual moving particles and detected different motion-type frequencies (Fig. 3G). The majority of particles assumed an active motion (52.36 ± 6.96 % for α>1.5) while we could also detect a small percentage of diffusive (5.04 ± 1.33 % for 0.9<α<1.1) and confined (8.86 ± 1.58 % for α<0.5) particles (Fig. 3H). As above, exogenous and endogenous pre-miRNAs appeared to behave similarly, since the computed α (α = 1.78 (endogenous) versus α = 1.79 (exogenous), Mann Whitney, p-value = 0.8797) (Fig. 3F) and α distribution (Fig. 3G,H) were not significantly different. Particle trajectory of each type of motion appeared to differ as illustrated by representative examples shown in Fig. 3I-K. Most trajectories analyzed displayed a perfect or near perfect fitting reflecting a single-mode behavior (Fig. 3I-K). These data together suggest that the majority of moving pre-miRNAs exhibit an active, directed motion along the axon. To gain insight into the added biological value of active transport, we computed the diffusion coefficient D (see Methods for details) for those particles moved only by diffusion (0.9<α<1.1). If we consider that the distance from *Xenopus laevis* RGC cell bodies to the tip of axons at this stage is 500 μm (Turner-Bridger et al., 2018), pre-miRNA puncta would take 20 days on average to reach the growth cone by diffusion while it takes merely two days for *Xenopus laevis* RGC axons to navigate to their main target (Holt and Harris, 1983). Collectively, these data suggest that the majority of anterogradely displaced pre-miRNAs are actively transported to promptly reach the growth cone.

### Pre-miRNAs are actively trafficked along microtubules, associated with vesicles

Next, we dissected the molecular mechanisms mediating pre-miRNA axonal transport. First, we explored whether pre-miR-181a-1 is trafficked along microtubules (MT), as axonal directed motion was previously shown to rely on this cytoskeletal component (Maday et al., 2014). We captured time-lapse movies before (-) or after (+) a 30 min incubation with 2.4 µM Nocodazole, a MT destabilizing drug, at a concentration sufficient to rapidly disrupt MT in *Xenopu*s RGC axons (Leung et al., 2018) (Fig. 4A). The drug did not affect the fluorescent signal *per se*, since particle fluorescence was unaltered (Fig. 4B). Kymograph analysis revealed a strong and significant reduction of the proportion of fast moving (> 0.5 μm/s) puncta (36.61 ± 2.08 % before versus 7.85 ± 5.10 % after treatment) and a significant increase in the proportion of stationary (< 0.2 μm/s) puncta (46.31 ± 2.09 % before versus 87.34 ± 7.83 % after treatment) (Fig. 4B,C). Furthermore, average velocities of all puncta significantly slowed down with the majority exhibiting no motion by the end of the treatment (median speed: 0.24 µm/s before to 0.02 µm/s after treatment) (Fig. 4D). Taken together, these results indicate that pre-miRNA transport along the axon shaft is mediated via microtubules.

**Figure 4:**
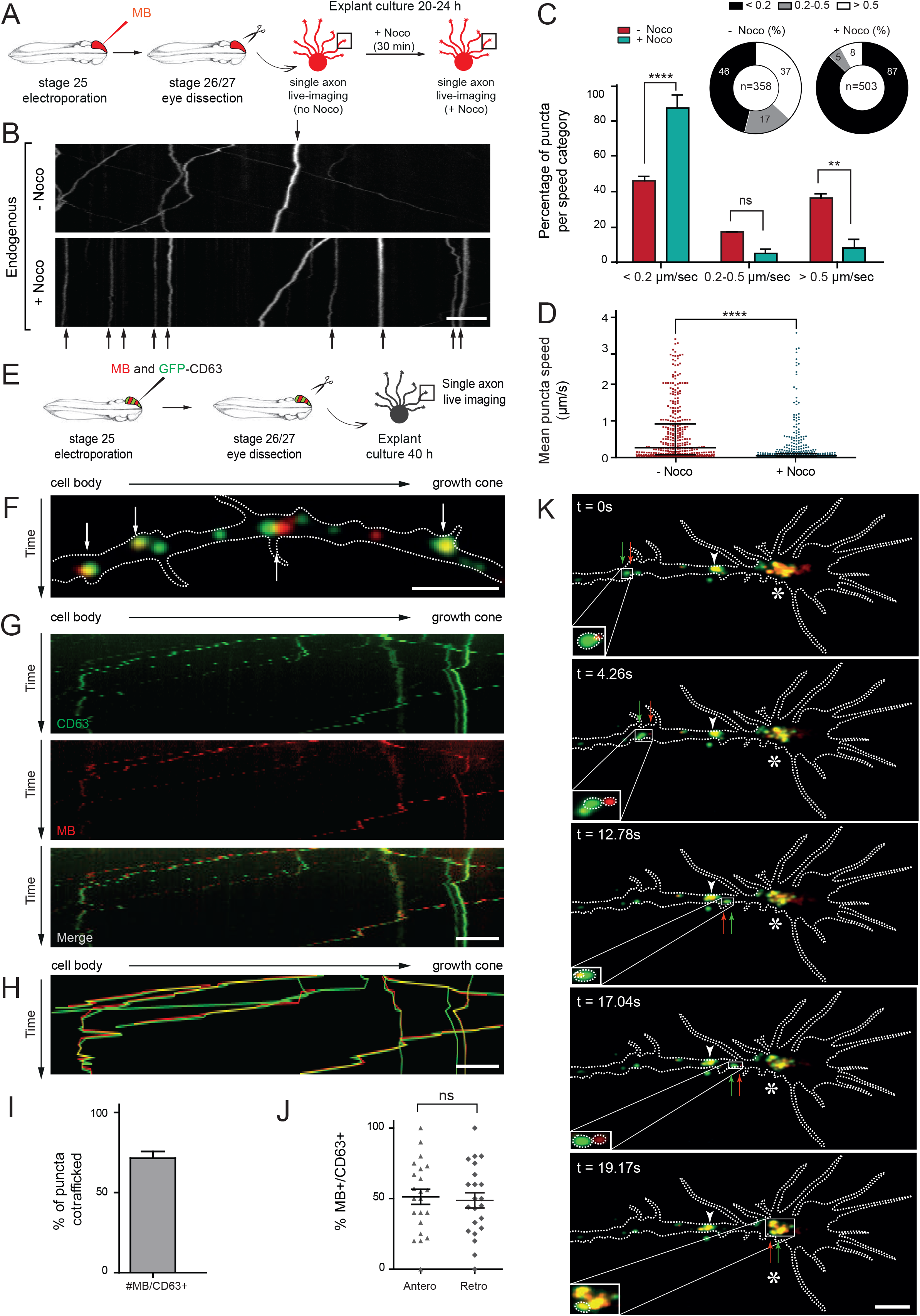
Pre-miR-181a-1 trafficking is vesicle mediated. **(A, E)** Schematic of the experimental paradigm. **(A)** 5 µM MB was electroporated. **(B)** Representative kymographs generated from the acquired time-lapse, before Nocodazole (top panel; - Noco) and 30 min after 2.4 μM Noco bath-application (bottom panel; + Noco). Stationary puncta in both panels are indicated with black arrows. **(C)** Frequency distribution (in percentage) and **(D)** mean MB punctum speed before (-) and after (+) Noco treatment. **(E)** 5 µM MB and 0.5 µg/ul pCS2-CD63-eGFP were co-electroporated. **(F)** Snapshot of a representative axon where MB-labeled pre-miR-181a-1 (red) and CD63-GFP-labeled vesicles (green) are co-trafficked (white arrows). **(G)** Representative kymographs generated from acquired time-lapse. **(H)** Composite kymograph shown in (G) where the individual traces where drawn and color coded. Yellow trajectories represent co-trafficked MB-labeled pre-miRNA (red) and CD63-GFP-labeled vesicle (green). **(I)** Frequency and **(J)** frequency distribution (in percentage) of MB-labeled pre-miR-181a-1 co-trafficked with CD63-GFP-positive vesicles. Each dot corresponds to % of MB+/CD63+ co-trafficked puncta within each axon (J). **(K)** Representative time-lapse depicting MB-labeled pre-miR-181a-1 (red arrow) and CD63-GFP-positive vesicle (green arrow) co-trafficked along the axon shaft to the growth cone (delineated with dashed white lines) wrist (white arrowhead) and central domain (white star). Values are mean ± SEM (C, I, J) or median with interquartile range (D). Statistics: ** p<0.01, **** p<0.0001. **(C, D)** n=3 independent experiments. Total number of analyzed puncta: 358 (- Noco) and 503 (+ Noco). **(C)** Two-way ANOVA followed by Tukey’s multiple comparison post-hoc test. **(D)** Data were not normally distributed (Shapiro-Wilk test), two-tailed Mann Whitney test. **(I)** Total number of counted puncta: 253 (MB+), 306 (CD63+). **(J)** Data were normally distributed (Shapiro-Wilk test), unpaired two-tailed t-test. Total number of puncta counted: 92 (anterograde), 78 (retrograde). **(I, J)** 22 axons, 5 independent experiments. Abbreviations: CD63, CD63-GFP; ns, not significant; MB, molecular beacon. Scale bars: 5 µm (B, F, G, H, K). *See also Figure S4*.

In neurons, mRNAs are packaged within ribonucleoparticles (RNPs) and trafficked along MT to distal neurites and back (Bauer et al., 2017). In agreement with this established means of transport, pre-miRNAs may be dynamically trafficked within RNPs, as recent data on dendrites suggest (Bicker et al., 2013). However, mature miRNAs, miRNA-repressible mRNAs and components of the miRNA processing machinery associate with late endosomes/multivesicular bodies and lysosomes (LE/Ly) in non-neuronal cells (Gibbings et al., 2009; Lee et al., 2009) and LE/Ly are detected in axons and growth cones (Falk et al., 2014; Konopacki et al., 2016). It is thus possible that pre-miRNAs adopt a non-canonical mode of transport associated with LE/Ly within the axon shaft. To explore this, we used CD63 as a LE/Ly marker, fused to GFP. CD63 is a transmembrane protein enriched in this compartment (Pols and Klumperman, 2009). We first examined whether CD63-GFP- and MB-labeled pre-miR-181a-1 puncta were co-trafficked within single stage 26/27 RGC axons following targeted eye electroporation (Fig. 4E). We detected that 71.41 ± 4.23 % of MB-positive puncta were co-transported with CD63-GFP-labeled vesicle-like focal puncta and an equal percentage of these moved anterogradely and retrogradely (Fig. 4F-J, Movie 3). Similar results were obtained when growth cones were cultured from older stage 37/38 embryos (#MB/CD63-GFP+: 73.65 ± 4.78 %) (Fig. S4A-G, Movie 4). These percentages are likely an underestimation, as endogenous unlabeled vesicles present in these axons may mask the extent of this co-traffick. These results indicate that CD63-positive vesicles contribute, to a large extent, to pre-miRNA axonal transport. Remarkably, the MB- and CD63-associated signals, while co-localizing, did not completely overlap (white arrows, Fig. 4F,S4B), indicating that pre-miRNAs may not reside inside vesicles, but may be tethered to them. Additionally, we investigated whether pre-miRNAs are transported to growth cones by hitchhiking on CD63-positive vesicles. We detected that numerous co-trafficked puncta reached the growth cones and appeared to stall within the organelle-rich central domain (Dent and Gertler, 2003) (star, Fig. 4K, Fig. S4G). We also observed a secondary storage point at the growth cone wrist, where MTs become bundled into dense parallel arrays (Bielas et al., 2007) in 70 % of axons analyzed (arrowhead, Fig. 4K, Fig. S4G). Taken together, these results suggest that pre-miRNAs are transported tethered to vesicles to the growth cone central domain where they are stored.

### Pre-miRNAs are processed locally in response to cues

Our data thus far indicate that miRNAs are actively delivered to growth cones, in which Dicer is also present, in their presumably inactive precursor form. Growth cones are sensory units that perceive and transduce chemotropic cues present in the environment. We wondered whether axonal pre-miRNAs could be locally processed into active mature miRNAs upon cue exposure, thereby contributing to growth cone turning. Although exogenous hairpin-less short double stranded RNA molecules designed to mimic endogenous small mature miRNAs (Ambion’s “Pre-miR™ miRNA Precursor Molecules”) are known to lead to the increase in mature miRNA in axons upon transfection (Aschrafi et al., 2008; Kar et al., 2013), nothing is known about local processing of *bona fide* pre-miRNAs.

To address this, we used isolated axons. We chose this approach over other widely used methods (Kim and Jung, 2015; Wang and Bao, 2017), namely compartmentalized chambers, Boyden chambers, or LCM axons, as these methods are not able to prevent communication between the axon and soma that remain physically connected. Isolated axons were prepared by carefully dislodging and manually removing the entire explant from the culture. In our hands, axonal and growth cone health is preserved with this approach, and growth cones remain responsive to cues (Fig. S5A). Furthermore, this approach yields pure axons that are devoid of the dendritic marker MAP2 (Fig. S5B) and are, thus, not contaminated by the somatodendritic compartment. We exposed isolated axons to well described repellent cues at concentrations that generate protein synthesis-dependent growth cone response (Bellon et al., 2017) (Fig. S5C) and we assessed whether any of these cues triggered pre-miRNA processing (Fig. 5D,E). For this, we quantified the levels of pre-miRNAs and miRNAs by RT-qPCR (Fig. 5A). Pre-miR-181a-1 and pre-miR-181a-2 are derived from two distinct primary transcripts and both give rise to miR-181a-5p and, respectively, to miR-181a-1-3p and miR-181a-2-3p (Fig. 5B-C). In *Xenopus*, the miR-181 family is composed of miR-181a (5p and 3p) and miR-181b, while miR-181c and d are unique to mammals (Kos et al., 2016; Wang et al., 2015). Following a Sema3A bath application to explants, we detected a significant increase in mature miR-181a-5p (+207.97 %), miR-181a-1-3p (+124.98 %) and miR-181a-2-3p (+194.21 %) (Fig. 5D). We further measured a significant decrease in pre-miR-181a-1 (−44.07 %) and pre-miR-181a-2 (−28.73 %) but not in pre-miR-182 (−1.94 %) (Fig. 5D). Additionally, Slit2 exposure did not significantly alter the levels of any of the tested miRNAs including miR-181a-5p (−1.26 %), miR-181a-1-3p (−8.82 %) and miR-181a-2-3p (+15.85 %) (Fig. 5E), and pre-miRNAs including pre-miR-181a-1 (−21.13 %), pre-miR-181a-2 (−20.22 %), pre-miR-182 (−25.45 %). Of interest, miR-181a-5p levels were far higher than miR-181a-1-3p and miR-181a-2-3p (24.30 and 10.11 fold difference respectively) following Sema3A exposure (Fig. S5D), suggesting that the 3p forms are rapidly degraded and unlikely to be functional. Taken together, these results suggest that Sema3A but not Slit2 triggers the processing of pre-miR-181a-1 and to some extent of pre-miR-181a-2 into mature miR-181a-5p, miR-181a-1-3p and miR-181a-2-3p. Since pre-miR-182 expression level is not altered, our results further indicate that specific cues induce the maturation of specific axonal pre-miRNAs. Exogenous pre-miRNA were recently demonstrated to be processed in dendrites in response to glutamate (Sambandan et al., 2017), pointing to the exciting possibility that cue-induced local miRNA maturation might be a key mechanism across different compartments.

**Figure 5:**
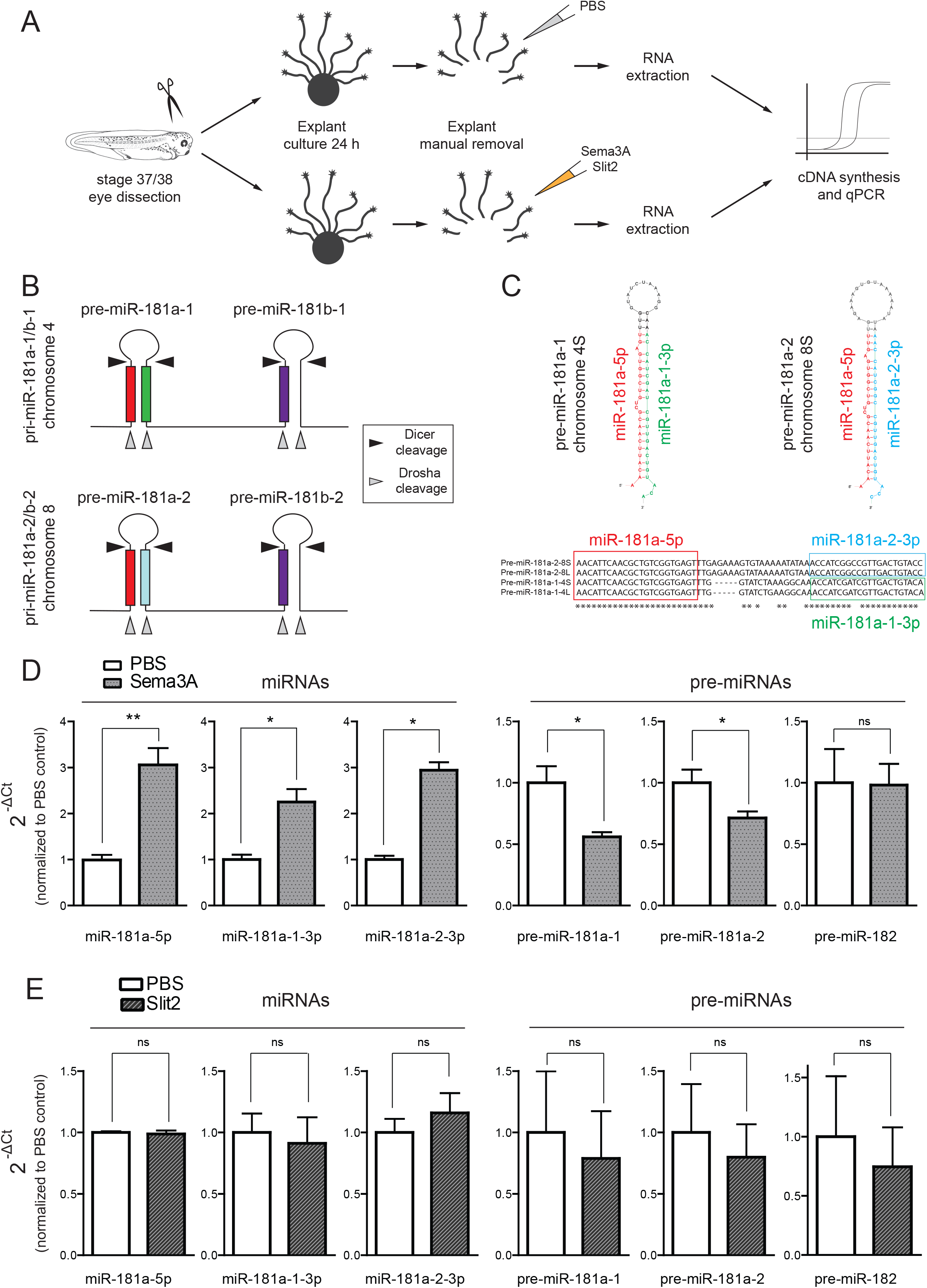
Pre-miRNAs are specifically processed locally upon cue stimulation. **(A)** Schematic representation of the experimental paradigm. Sema3A and Slit2: 200 ng/mL. **(B)** Schematic of pri-miR-181a-1/b-1 on chromosome 4 and of -181a-2/b-2 on chromosome 8. Small gray arrowheads indicate Drosha cleavage sites and the black ones Dicer cleavage sites. Colored boxes show the position of mature miRNAs. **(C)** Predicted secondary structure (Mfold v3.6) and multiple alignment (MUSCLE v3.8) of the two precursors isoforms, pre-miR-181a-1 (chromosome 4S) and pre-miR-181a-2 (chromosome 8S). **(D, E)** Quantification of miRNA and pre-miRNA expression levels using the 2^^(-ΔCt)^ method and U6 as normalizer, upon Sema3A (D) or Slit-2 (E) stimulation. Data are normalized to PBS control. Values are mean ± SEM. Statistics: * p<0.05, ** p<0.01. Data were not normally distributed (Shapiro-Wilk test). Two-tailed Mann Whitney test, n=3-4 independent experiment. Abbreviations: ns, not significant. *See also Figure S5*.

### Newly synthesized miRNAs are important for growth cone steering ***ex vivo***

We next explored whether newly synthesized miRNAs (NSmiRNAs) are important players in modulating growth cone behavior. We reasoned that if they are indeed crucial, then blocking their production should impair growth cone responsiveness to cues. We thus blocked miRNA biogenesis by preventing Dicer-mediated cleavage with a mix of two morpholinos (MOs) complementary to 5p (MOs-5p) or 3p (MOs-3p) Dicer cleavage site of both pre-miR-181a-1 and -2 (Fig. 6A,B). First, we established that these MOs successfully impair pre-miRNA processing *in vivo*. For this, we microinjected MOs at the 8 cell stage into cells fated to become CNS, and measured miRNA levels by qPCR from stage 40 retinal extract (Fig. S6A). As expected, MOs-5p and MOs-3p both lead to the strong reduction of mature miR-181a-5p (−98.41 % [MOs-5p] and -87.36 % [MOs-3p]), miR-181a-1-3p (−63.20 % [MOs-5p] and -83.02 % [MOs-3p]) and miR-181a-2-3p (−92.89 % [MOs-5p] and -96.55 % [MOs-3p]) (Fig. S6B,C), confirming that these MOs block miRNA biogenesis. Next, we assessed whether these MOs prevent Sema3A-induced pre-miR-181a-1 and pre-miR-181a-2 processing from pure isolated axons *ex vivo* (Fig. S6D). Stage 37/38 isolated axons were transfected with MOs-5p or -3p, and subsequently bathed with Sema3A or PBS for 10 min (Fig. S6D). The level of the mature miRNA stemming from the strand opposite to that complementary to MO was measured by TaqMan PCR (Fig. S6D). As expected, Sema3A induced a significant increase in axonal miR-181a-1-3p and -5p levels in co-MO transfected axons (+103.59 [3p]; +64.23 [5p]) indicative of pre-miRNA processing but not in MOs-5p and -3p transfected axons (+1.42 [3p]; +3.83 [5p]) (Fig. S6E,F). This indicates that the MOs block Dicer-induced pre-miRNA cleavage in axons and also confirms the specificity of Sema3A-induced pre-miR-181a-1 and pre-miR-181a-2 processing observed earlier.

We then examined whether blocking pre-miRNA processing, as above, impairs growth cone responsiveness to Sema3A. We employed a similar experimental paradigm except that this time we assessed growth cone behavior by collapse assay (Fig. 6C). Sema3A induced growth cone collapse in isolated axons transfected with co-MO (Fig. 6D: 26.37 ± 2.76 % [PBS]; 56.97 ± 2.03 % [Sema3A]; Fig. 6E: 29.17 ± 1.43 % [PBS]; 61.13 ± 2.76 % [Sema3A] collapsed growth cones). In contrast, a significant reduction in Sema3A-induced collapse was observed in axons transfected with MOs-5p or -3p (MOs-5p (Fig. 6D): 33.89 ± 3.73 % [PBS]; 36.71 ± 0.53 % [Sema3A]; MOs-3p (Fig. 6E): 33.12 ± 2.67 % [PBS]; 46.04 ± 0.61 % [Sema3A] collapsed growth cones). These results reveal that pre-miR-181a-1 and pre-miR-181a-2 processing is required for growth cone responsiveness to Sema3A. They further suggest that NSmiRNAs impinge on the Sema3A signaling pathway by targeting transcripts important for Sema3A-mediated growth cone turning.

**Figure 6:**
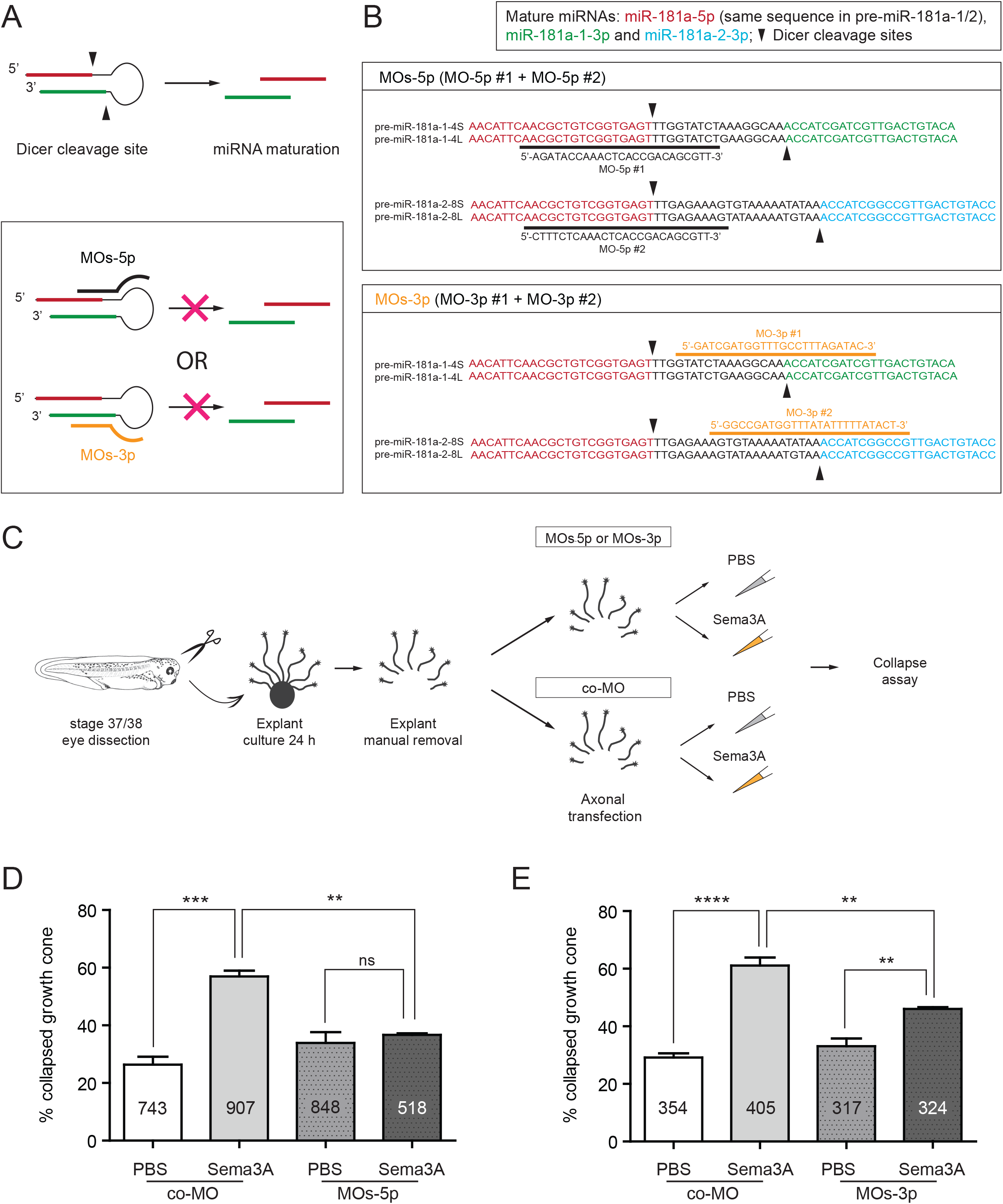
Pre-miRNAs processing mediates growth cone responsiveness to Sema3A. **(A, top panel)** Schematic of pre-miRNA maturation upon Dicer cleavage. **(A, bottom panel)** Schematic of MOs-5p and MOs-3p targeting 5’ and 3’ pre-miRNA Dicer cleavage site respectively. **(B)** Exact MO targeting region within pre-miR-181a-1 and -2 derived from chromosomes 8, and 4, long (L) or short (S). Small black arrowheads indicate Dicer cleavage sites. **(C)** Schematic representation of the experimental paradigm. **(D, E)** Frequency (in percentage) of collapsed growth cones from stage 37/38 embryos, following a 10 min (200 ng/mL) Sema3A bath application. Mix 2 µM MOs-5p (D) or 2 µM mix MOs-3p (E) were used. Total number of counted growth cones is reported in the column. Values are mean ± SEM. Statistics: ** p<0.01, *** p<0.001 and **** p<0.0001. **(D, E)** Two-way ANOVA followed by Tukey’s multiple comparison post-hoc test, n=3 independent experiments. Abbreviations: ns, not significant. *See also Figure S6*.

### Mature miRNAs are important for axon targeting ***in vivo***

As NSmiRNAs are essential for growth cone behavior *ex vivo*, we next assessed whether NSmiRNAs are also important for growth cone turning *in vivo*. To address this, we use a miR-181-MO cocktail to block formation and function of mature miRNAs. We electroporated miR-181-MO directly into stage 26 developing eyes, bypassing all earlier developmental stages, and examined axon projections at stage 40, when pioneer axons have reached the optic tectum (Holt and Harris, 1983) (Fig. 7A). While long-range guidance was unaffected, a subset of axons strayed within the optic tectum following aberrant trajectories (Fig. 7B, arrows) as revealed by quantification (Fig. 7C). Though a few straying axons are always observed within the wild type tectum, this number was significantly increased upon depletion of miR-181 (8.6 ± 1.01 % [co-MO]; 21.8 ± 2.65 % [miR-181-MO]) (Fig. 7C). To assess the specificity of miR-181-MO, we used miRNA mimics, double stranded miRNAs that are able to restore miRNA expression as confirmed by *in situ* hybridization (ISH) (Fig. S6A-C). Serial electroporation of miR-181-MO and of miRNA mimics into stage 26 embryos rescued the phenotype observed at stage 40 (4.05 ± 1.89 % [co-MO+control mimics]; 17.46 ± 3.19 % [miR-181-MO + control mimics]; 5.09 ± 2.13 % [miR-181-MO + miR-181 mimics]) (Fig. 7D,E). This confirms the specificity of the miR-181-MO. Taken together, these results suggest that mature miRNAs are important for axon targeting *in vivo*. Since the onset of Dicer temporal expression occurs when axons are reaching their target (Fig. 1D), these results overall suggest a restricted role of NSmiRNAs during this critical decision period. They further confirm that miRNAs are crucial players in axon targeting (Baudet et al., 2013; Iyer et al., 2014).

**Figure 7:**
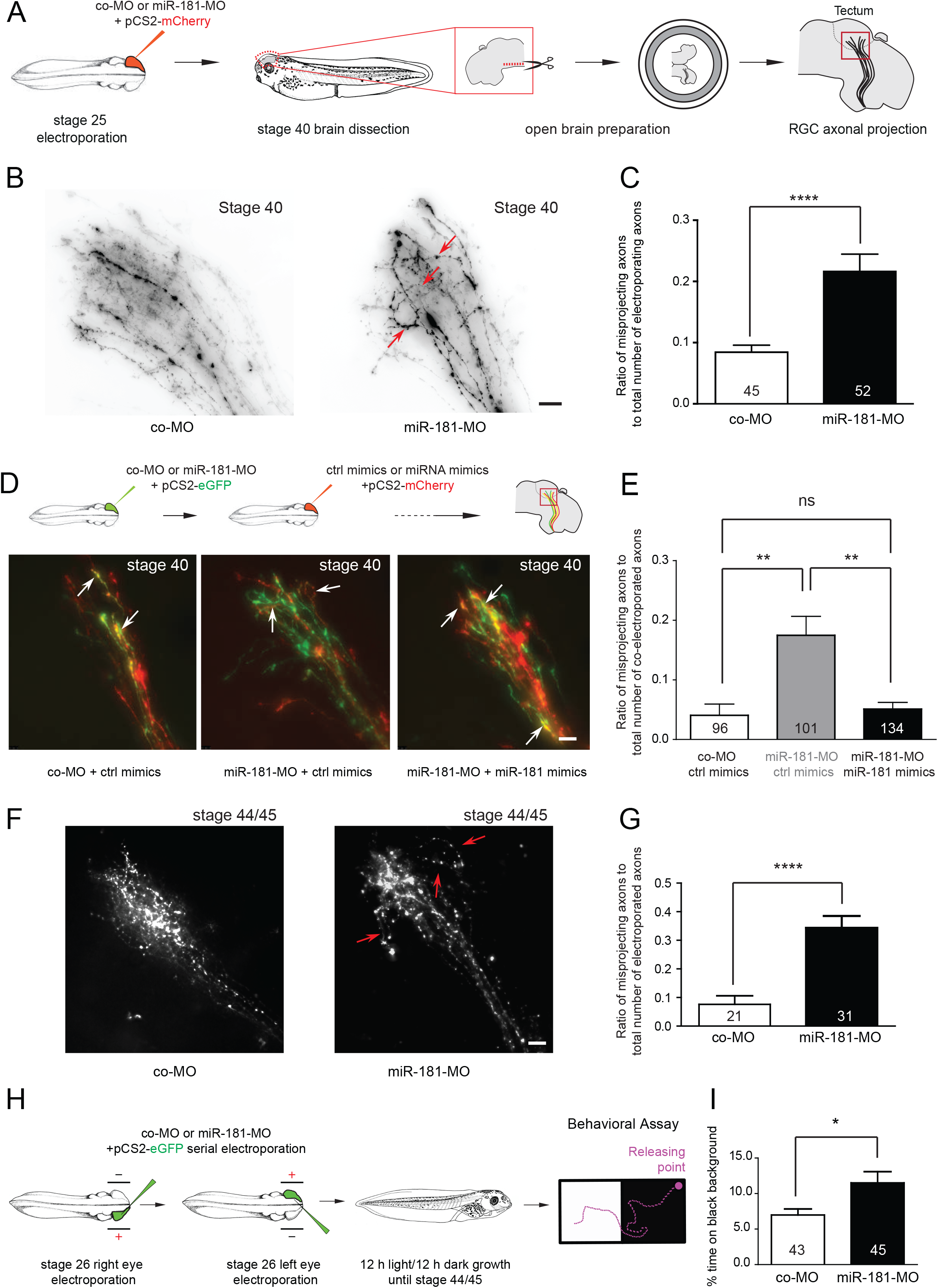
miR-181 loss-of-function leads to aberrant axon targeting and impaired vision. **(A, D top, H)** Schematic representation of the experimental paradigm. Concentrations used for electroporation: 0.5 μg/μl pCS2+mCherry or pCS2+eGFP plasmids; 250 μM miR-181-MO cocktail (62.5 μM pre-miR-181a-1-MO plus 62.5 μM pre-miR-181a-2-MO plus 125 μM miR-181b-MO) or 250 μM control MO cocktail (co-MO; 125 μM custom control plus 125 μM standard control); 50 μM miR-181 (25 μM miR-181a-, plus 25 μM miR-181b-mimic) or 50 μM control mimics. miR-181 MO cocktail was used to block the processing of pre-miR-181a-1 and pre-miR-181a-2 and thus the function of miR-181a-5p, -1-3p and -2-3p. Since miR-181a-5p and miR-181b share the same seed and differ by only 4 nucleotides, we also used a MO to block miR-181b function to avoid any functional redundancy. **(B, D bottom, F)** Representative images of RGC axons within the optic tectum. **(B, F)** A subset of aberrantly projecting axons are indicated (red arrows). Axons were considered as straying when they abnormally looped, twisted or bent within the tectum. **(D)** White arrows indicate axons targeted both by MO (green) and miRNA mimics (red). **(C, E, G)** Quantification of misprojecting axons. The number reported on the bars are the total number of analyzed brains **(C, G)** or co-electroporated axons **(E)**. Note how miR-181 mimics rescued aberrant misprojection of morphant axons *in vivo*. **(I)** Frequency (in percentage) of the amount of time embryos spent on black background. Numbers on the columns are the total number of analyzed embryos. Values are mean ± SEM. Statistics: * p<0.05, ** p<0.01 and **** p<0.001. **(C, G, I)** Data were not normally distributed (Shapiro-Wilk test). Two-tailed Mann Whitney test, n=4 independent experiments. **(E)** Data were normally distributed (Shapiro-Wilk test). One-way ANOVA followed by Tukey’s multiple comparison post-hoc test, n=4 independent experiments. Abbreviations: co-MO, control morpholino; ns, not significant. Scale bars: 20 µm (B, D, F). *See also Figure S7*.

By stage 44/45, a similar subset of miR-181-MO electroporated axons projected aberrantly within the tectum (7.62 ± 2.98 % [co-MO]; 34.35 ± 4.17 % [miR-181-MO]) (arrows, Fig. 7F,G) suggesting that straying axons are not eliminated at later stages due to lack of trophic support (Yamaguchi and Miura, 2015) and that this phenotype is therefore not transient. Finally, we evaluated whether these straying axons impact vision by performing a behavioral assay which measures visual discrimination between white and black backgrounds (Viczian and Zuber, 2014) (Fig. 7H). Contrary to blind embryos, WT embryos normally seek light-colored environment, and therefore spend overall little time on a black background (Fig. S7D). Following serial miR-181-MO electroporation into right and left retinas at stage 26, stage 44/45 embryos displayed a significant decrease in the preference for white background compared to co-MO controls (6.99 ± 0.84 % [co-MO]; 11.51 ± 1.57 % [miR-181-MO]) (Fig. 7I). The miR-181 family is therefore important to ensure accurate targeting of RGC axons within the tectum and subsequently for fully functional vision.

### NSmiRNAs silence locally translated transcript

To explore the molecular mechanisms through which NSmiRNAs act, we reasoned that NSmiRNAs could either 1) silence translationally active transcripts acting as a switch, 2) prevent translational onset of axonal mRNAs as a fail-safe mechanism to avoid spurious translation, or 3) degrade unneeded or unwanted transcripts to promote growth cone turning. We ruled out the latter possibility, since the combined processes of miRNA biogenesis and mRNA degradation are unlikely to occur within the time frame of cue stimulation (5-10 min). We thus explored the effect of NSmiRNAs on local translation.

We first identified putative direct targets of NSmiRNA miR-181a-5p, the most abundant axonal mature miRNAs derived from pre-miR-181a-1 and pre-miR-181a-2 (Fig. S5D). Following total RNA-seq from stage 37/38 isolated axons, we shortlisted axonally expressed mRNAs that complied with the following criteria (see Methods for details): 1) a minimum 50 bp 3’UTR length, 2) at least one miR-181 miRNA responsive elements (MRE), 3) among the top 20 % of predicted targets (based on TargetScan’s context score), and 4) predicted miR-181 targets in either human or mouse (Table 1). Since blocking NSmiRNA biogenesis impaired Sema3A-induced collapse (Fig. 6D,E), we inferred that under normal conditions, the NSmiRNA-induced silencing of candidate mRNAs would assist the collapse response. It could do so by supporting mechanisms involved in repulsive turning such as by impairing cytoskeleton polymerization, impairing cell adhesion to laminin and/or by enabling Sema3A signaling. We selected one representative miR-181 target candidate reflecting each of these three possibilities. First, we chose to focus on tubulin beta 3 class III (TUBB3), a microtubule beta isotype needed for proper axon guidance and targeting (Poirier et al., 2010; Tischfield et al., 2010), and ranked second among the putative targets of the axon guidance Reactome pathway (R-HSA-422475) (Table 1). We also selected Thrombospondin 1 (THBS1), an adhesive glycoprotein mediating the interaction between cells and the extracellular matrix (Resovi et al., 2014) and the top ranked target among the members of the integrin cell surface interaction Reactome pathway (R-HSA-216083) (Table 1). Integrins are transmembrane receptors mediating axonal adhesion to laminin (Yamada and Sekiguchi, 2015). Finally, we selected amyloid beta precursor protein (APP) known to prevent Sema3A-induced collapse (Magdesian et al., 2011) with clinical relevance in neurodegenerative diseases (Roher et al., 2017).

We assessed whether local translation of these three candidate mRNAs could be regulated by NSmiRNAs using FRAP (fluorescence recovery after photobleaching) of a fast folding and fast bleaching translational reporter, Venus, carrying the 3’UTR of genes of interest (Ströhl et al., 2017; Wong et al., 2017) (Fig. S8A). We first tested that FRAP can be successfully used to measure LPS of a well-established axonal mRNA, *β*-Actin (ACTB). We conducted FRAP on *ex vivo* RGC growth cones following the targeted eye electroporation of Venus-ACTB-3’UTR construct and mRFP as a general cell marker (Fig. 8A). We observed a rapid 38.28 ± 5.98 % fluorescence recovery at 10 min in growth cones expressing Venus-ACTB-3’UTR and a significant lower recovery 22.75 ± 2.61 % when 100 μM cycloheximide (CHX), a translational blocker, was applied (Fig. S8B,C). These results show that FRAP of Venus-3’UTR construct can be used to analyze local regulation of transcripts in growth cones.

**Figure 8:**
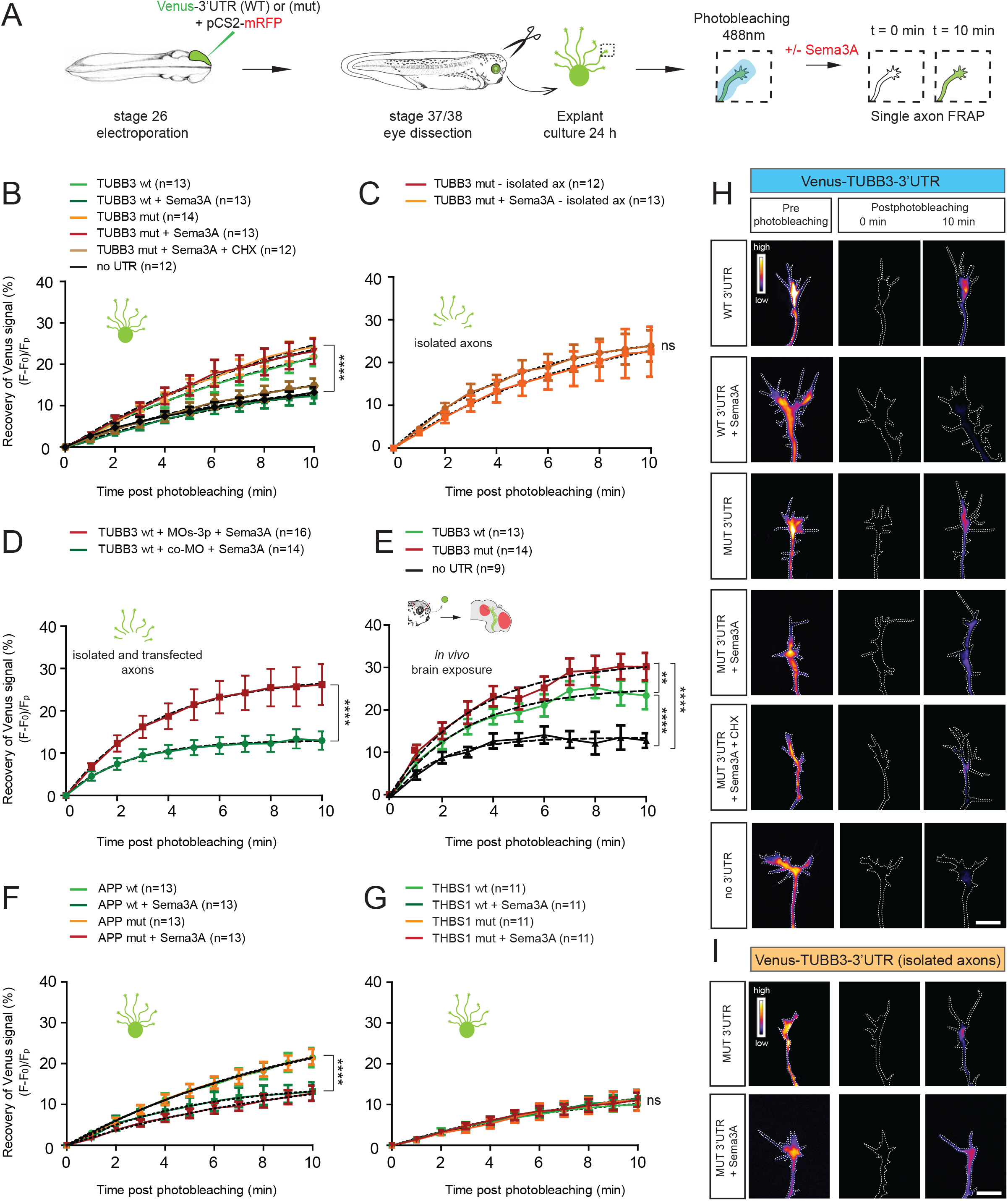
FRAP: *ex vivo* and *in vivo* 3’UTR TUBB3 regulation by miR-181-5p. **(A)** Schematic representation of the experimental paradigm. Plasmid concentrations used: 1 μg/μl of pCS2+mRFP; 0.7 μg/μl of pCS2+Venus-no 3’UTR/pCS2+Venus-3’UTR of interest. **(B-G)** Quantification (in percentage) of the axonal fluorescence recovery after photobleaching (FRAP) of different Venus-3’UTR constructs *ex vivo* and *in vivo. Ex vivo*, whole explants (B, F, G), isolated axons (C) or isolated transfected axons (D) were used for FRAP analysis. *In vivo* (E), whole embryos with exposed brain were used. The electroporated eye was removed prior to mounting the embryo to eliminate somatic contribution. The red area on the brain schematic indicate Sema3A expressing territories. **(B)** 200 ng/ml Sema3A was bath applied with or without 100 µM cycloheximide (CHX, a translational blocker). **(D)** Axons were transfected with 2 µM control morpholino (co-MO) or 2 µM MOs-3p. **(H, I)** Representative growth cones depicting Venus fluorescence intensity as a heatmap. Values are mean ±SEM. Statistics: ** p<0.01 and **** p<0.0001. **(B-G)** Numbers of single axons analyzed are reported between brackets. n=5 (B), n=4 (C), n=3 (D-G) independent experiments. Dashed black lines represent least-square fits to a single-exponential decay equation. Exact p-value of all FRAP experiments and row statistics are reported in Table 4. Abbreviations: ns, not significant; wt, wild type; mut, miR-181-5p responsive elements mutated; CHX, cyclohexamide; TUBB3, tubulin beta 3 class III; APP, Amyloid beta precursor protein; THBS1, Thrombospondin 1. Scale bars: 10 µm (H, I). *See also Figure S8*.

We then sought to examine the *de novo* synthesis of TUBB3 using the Venus-TUBB3-3’UTR construct (Fig. S8A). Growth cones expressing Venus alone (no 3’UTR) displayed a minimal amount of recovery within 10 min post-photobleaching (13.24 ± 1.22 %). This corresponds to diffusion of Venus from adjacent, non-bleached regions to the bleached growth cone (Wong et al., 2017). By contrast, Venus-TUBB3-3’UTR expressing growth cones exhibited rapid recovery within this 10 min-time frame (21.85 ± 2.30 %) (Fig. 8B,H). Sema3A exposure suppressed the fluorescence recovery in Venus-TUBB3-3’UTR growth cones to levels similar to Venus alone (12.32 ± 1.81 % at 10 min) (Fig. 8B,H). These data suggest that TUBB3 is rapidly synthesized in growth cones and that Sema3A prevents local translation of TUBB3.

Next, we tested the role of mature miRNAs in mediating Sema3A-induced repression of TUBB3. To this end, we mutated the two miR-181a-5p MREs within TUBB3 3’UTR(mut) to decouple Sema3A-induced miRNA maturation from mRNA binding and silencing (Fig. S8A). Mutating TUBB3 3’UTR did not affect fluorescence recovery in basal conditions (24.14 ± 2.23 % at 10 min) compared to WT (21.85 ± 2.30 % at 10 min) indicating that mature miRNAs do not regulate the constitutive TUBB3 expression in distal axons (Fig. 8B,H). Furthermore, upon Sema3A exposure, fluorescence recovery in growth cones expressing Venus-TUBB3-3’UTR(mut) (23.15 ± 3.08 % at 10 min) was significantly higher than that of growth cones expressing Venus-TUBB3-3’UTR(WT) (12.32 ± 1.81 % at 10 min), and similar to that observed for Venus-TUBB3-3’UTR(mut) without Sema3A (24.14 ± 2.23 % at 10 min) (Fig. 8B,H). This indicates that Sema3A triggers mature miRNA activity, which in turn represses TUBB3. CHX abolished the recovery of Venus-TUBB3-3’UTR(mut) expressing growth cones bathed with Sema3A (14.77 ± 1.79 % at 10 min) (Fig. 8B,H), indicating that TUBB3(mut) is still translated even in the presence of Sema3A and, therefore, that repression does not occur without mature miRNAs. Similar results were obtained when isolated axons were used (Venus-TUBB3-3’UTR(mut) [no Sema3A]: 23.90 ± 3.66 %; Venus-TUBB3-3’UTR(mut) [+Sema3A]: 22.52 ± 5.83 % at 10 min) (Fig. 8C,I). Taken together, these results reveal that repression of TUBB3 translation is mediated by Sema3A-activated, mature miRNAs locally within axons. To confirm that these activated, mature miRNAs are in fact NSmiRNAs, we interfered with the biogenesis of NSmiRNAs prior to FRAP. For this, we transfected isolated axons with co-MO or MOs-3p that block the processing of pre-miRNAs (Fig. S8D). Upon Sema3A exposure, growth cones expressing Venus-TUBB3-3’UTR(WT) exhibited a significantly higher recovery when transfected with MOs-3p (26.20 ± 4.82 % at 10 min) compared to those transfected with co-MO (12.98 ± 2.17 % at 10 min) (Fig. 8D). This indicates that NSmiRNAs mediate Sema3A-induced translational silencing of TUBB3 locally within axons.

We then investigated whether mature miRNAs control TUBB3 expression in RGC axons *in vivo*. For this, we expressed Venus-TUBB3-3’UTR (WT or mut) and mRFP in RGC by retinal electroporation and performed *in vivo* FRAP on RGC distal axons within the tectum in the vicinity of Sema3A expressing territories (Fig. S8E). The electroporated eye was removed to avoid diffusion of confounding soma-derived, Venus-tagged proteins into the axons (Fig. S8E). Growth cones expressing Venus alone displayed low level of signal recovery (15.07 ± 2.36 % at 10 min) following photobleaching. By contrast, growth cones expressing Venus-TUBB3-3’UTR(WT) displayed a rapid fluorescence recovery reaching 24.13 ± 2.96 % at 10 min. This indicates that TUBB3 is locally translated *in vivo* within the RGC targeting region. When growth cones expressed Venus-TUBB3-3’UTR(mut) instead, fluorescence recovery was significantly increased (30.72 ± 2.23 % at 10 min) (Fig. 8E). This further indicates that mature miR-181a-5p normally represses and fine tunes TUBB3 local translation in growth cones near sources of tectal Sema3A *in vivo* to promote accurate navigation within the target area. It further indicates that the observed impaired targeting in the absence of functional miRNAs (Fig. 7) might be attributed to excessive levels of TUBB3 in RGC axons at the tectum.

Finally, we assessed whether Sema3A-induced NSmiRNAs modulate the translation of APP and THBS1 (Fig. S8A). Fluorescence from Venus-APP-3’UTR expressing axons recovered following photobleaching (21.47 ± 2.35 % at 10 min), and this recovery was significantly impaired when growth cones were exposed to Sema3A (13.20 ± 2.25 % at 10 min) (Fig. 8F). This indicates that APP is locally translated in growth cones, and APP LPS is repressed by Sema3A. Since APP interferes with Sema3A-induced growth cone collapse (Magdesian et al., 2011), Sema3A may increase growth cone sensitivity to itself by controlling APP expression level. We, next, tested whether mature miRNAs could be a key mediator in Sema3A-mediated repression of APP by mutating MRE as above (Fig. S8A). Venus-APP-3’UTR(mut) expressing axons exhibited a fluorescence recovery (12.74 ± 1.87 % at 10 min) similar to that of WT 3’UTR (13.20 ± 2.25 % at 10 min) (Fig. 8F), suggesting that mature miR-181a-5p does not contribute to mediate Sema3A-induced silencing. Furthermore, Venus-THBS1-3’UTR(WT) expressing axons displayed similar recovery (9.99 ± 1.40 % at 10 min) compared to Sema3A-exposed WT axons (11.54 ± 2.11 % at 10 min) or when the three miR-181a-5p MREs within THBS1-3’UTR were mutated (mut) in the presence (11.17 ± 1.76 % at 10 min) or absence of Sema3A (11.05 ± 2.53 % at 10 min) (Fig. 8G). This suggests that THBS1 is not locally translated in axons. Collectively, this indicates that not all predicted miR-181 targets are regulated by mature miRNAs within RGC axons.

Taken together, these results demonstrate that Sema3A inhibits basal translation of key molecules. They further reveal that NSmiRNAs are a major component of the Sema3A signaling pathway that are required for the repression of specific, translationally active transcripts within growth cones, thereby acting as a switch.

## Discussion

In this study we show that pre-miR-181a-1 is actively transported, tethered to CD63-positive vesicles along axonal microtubules to the growth cone central domain using a novel approach based on MB. Sema3A bath application of isolated axons leads to the processing of pre-miR-181a-1 and pre-miR-181a-2 into mature miRNAs in a cue and pre-miRNA-specific manner. These mature miRNAs are important for growth cone collapse *ex vivo*, axon guidance *in vivo* and necessary for a normally functioning visual system. Mechanistically, cue-generated NSmiRNAs silence the basal translation of a specific transcript, TUBB3, at the growth cone *ex vivo* and *in vivo*. Collectively, our findings are consistent with a model in which pre-miRNAs are delocalized to and stored within growth cones in an inactive form. Upon cue exposure, they are rapidly processed into active miRNAs to inhibit the basal local translation of transcripts, thereby ensuring accurate axon trajectories.

### miRNAs are transported to the growth cone as inactive precursors via LE/Ly-like vesicles and specifically processed locally on demand

Transport of mRNA as a mean to delocalize genetic material is well described. While emerging evidence strongly suggest that ncRNAs and in particular miRNAs are differentially distributed in compartments (Iyer et al., 2014; Kye et al., 2007; Lugli et al., 2008; Natera-Naranjo et al., 2010), mechanisms leading to the compartmentalization of ncRNAs are largely unexplored. It is technically very challenging to study transport of endogenous miRNAs due to their short length, making classic tools such as the long MS2 tagging ineffective. To investigate mechanisms of miRNA dynamics, we have adapted MB, a technique already employed by others to track miRNA localization, although not their transport dynamics [e.g. (Földes-Papp et al., 2009)], and mRNA trafficking in living cells including in *Xenopus* RGC axons (Turner-Bridger et al., 2018). The series of data we provide here collectively suggests that the MB we designed specifically detects pre-miRNAs in axons. First, in RGC axons the MB is complementary only to pre-miR-181a-1 as shown through combined *Xenopus* genome blast and axonal RNA-seq analysis (see Method and Table 2). Second, the MB recognizes and binds to exogenous pre-miRNA *in vitro* as revealed by the thermal denaturation assay. Third, MB and exogenous pre-miRNA co-localize by at least 71 % within axons *ex vivo*. This level of co-localization appears particularly high considering that MB and pre-miRNA may not be delivered to axons with similar copy numbers when using serial electroporation, and that the MB are likely to also detect endogenous pre-miRNAs besides the exogenous ones. Fourth, pre-miRNA abundance detected by qPCR and number of MB-positive puncta in axons change to a virtually equal extent (56 % and 58 % decrease, respectively) in response to impairment of primary miRNA processing (Fig. 2 E, F). Fifth, MB and pre-miRNA trafficking dynamics are highly similar. MBs are thus a powerful new tool to specifically track pre-miRNA transport.

Recent studies have provided great insight into axonal mRNA trafficking (Leung et al., 2018; Turner-Bridger et al., 2018). Many components of pre-miRNA dynamics that we report here resemble those of mRNA in axons. We document similar bidirectional trafficking (Alami et al., 2014; Leung et al., 2018; Turner-Bridger et al., 2018) and faster anterograde than retrograde transport (Turner-Bridger et al., 2018). Maximal velocities and average speed of active particles (~0.8-1.1 μm/s (Turner-Bridger et al., 2018) vs ~1 μm/s (0.972 ± 0.08 μm/s (endo); 1.064 ± 0.08 μm/s (exo) [mean speed ± SEM] our work) are also comparable (Leung et al., 2018; Turner-Bridger et al., 2018). In contrast to our study, the dominant mRNA trajectories previously measured (Turner-Bridger et al., 2018) were confined and diffusive, with only a small proportion being directed, while the majority of particles we detected adopted an overall active and directed trajectory. This disparity in motion type frequencies can have a technical explanation, or reflect actual biological differences. We employed an MSD analysis of overall trajectory limited to moving particles, over a short temporal frame, while Turner-Bridger and collaborators used a bespoke analysis pipeline of segmental trajectories of all particles (Turner-Bridger et al., 2018). Alternatively, there could be *bona fide* biological differences in transport dynamics between mRNAs and pre-miRNAs. In axons mRNA can be reused for multiple rounds of translation while pre-miRNA cannot be reutilized, as they are processed. It is thus tempting to speculate that a constant supply of fresh pre-miRNAs may therefore be required to replenish the growth cone pre-miRNA storage. Since pre-miRNA diffusion to the growth cone would, on average, take 20 days, rapid active transport would be required to match the ever-changing demands of fast elongating axons.

Furthermore, we provide evidence demonstrating that pre-miRNA molecules are transported with vesicles. What is the nature of these vesicles? Several lines of evidence support the notion that pre-miRNAs are trafficked and stored in close association with LE/Ly. First, the vast majority (71-74 %, depending on the stage under study) of pre-miRNA puncta are co-trafficked along axons coupled to CD63-GFP-positive vesicles. CD63 is a small integral protein belonging to the tetraspanin superfamily (Charrin et al., 2014). While CD63 is found in the exocytic pathway and on the cell surface, like most transmembrane proteins, it is predominantly detected within LE/Ly (Pols and Klumperman, 2009). Second, markers of LE/Ly are chiefly located at the central domain within the growth cone, similarly to where pre-miRNA and CD63-GFP positive vesicles are detected. In contrast, markers for early and recycling endosomes are also found within the growth cone peripheral domain and filopodia in embryonic *Xenopus* RGC (Falk et al., 2014; Konopacki et al., 2016) where pre-miRNA / CD63-GFP were rarely observed (Fig. 3B,4K; Fig. S4G). Third, pre-miRNAs were detected in close proximity to CD63-positive vesicles. miRNAs, components of miRISC such as Ago2 and GW182, and miRNA-repressible mRNA associate with LE/Ly membranes in non-neuronal cells (Gibbings et al., 2009; Lee et al., 2009). GW-182, for example is juxtaposed to the outer limiting membrane of LE/Ly within the cytosol (Lee et al., 2009). Overall, our data suggest that pre-miRNAs are transported, tethered to the outer membrane of LE/Ly, to growth cones for subsequent storage. Pre-miRNAs are therefore not destined to be shipped to the extracellular milieu via exosomes but to act within axons.

Our results contrast with a previous report that has documented an association of exogenous pre-miRNAs (pre-miR-338 in rat SCGs) with mitochondria in axons (Vargas et al., 2016). Although we cannot rule out that pre-miR-181a-1 is linked to this organelle in small percentage or in addition to LE/Ly, this difference might be attributed to several factors including the type of miRNA and/or neuron under study. In contrast to Kaplan’s group which investigated PNS axons, we examine here CNS axons. Intriguingly, non-canonical hitchhiking onto membrane-bound vehicles such as ER, early endosomes, COPI and secretory vesicles has been described for a small subset of cargoes to achieve subcellular motility (Salogiannis and Reck-Peterson, 2017). In particular, mRNAs in fungi were found to translocate docked to cytoplasmic surface endosomes (Baumann et al., 2012). This suggests that the pre-miRNA mode of trafficking belongs to an ancient and evolutionary conserved transport system that spans across species and subcellular compartments.

Upon translocation to distal axons, we uncover that pre-miRNAs are stored within growth cones and processed into mature, active miRNAs on demand. Delocalizing miRNA biogenesis would present several key advantages. First, sequences within the pre-miRNA, such as the loop region, would allow the evolutionary acquisition of distinct and putative targeting motifs (Bicker et al., 2013; Smalheiser, 2008) which would otherwise not easily fit within the much shorter 22nt mature miRNA sequence. These motifs would subsequently aid the translocation of pre-miRNAs to specific subcellular compartments. Second, the transport of inactive precursors would avoid spurious activity along the transport route until the proper processing machinery – and signal – is encountered at the growth cone. This would constitute a fail-safe mechanism to compartmentalize signaling events at the right time, i.e. upon cue exposure, and at the right place, i.e. not only within the growth cone but perhaps to growth cone subdomains closest to the cue. Finally, pre-miRNAs would be readily available for immediate use by the growth cone on demand, contrary to mature miRNAs which would need to be transported from the soma to the distal axon upon cue-mediated activation. Overall, local processing of miRNAs into NSmiRNAs would be beneficial to the cell. Impaired pre-miRNA trafficking and concomitant local action of NSmiRNAs may be a hitherto overlooked etiological factor of neurodegenerative diseases.

### Basal local protein translation of TUBB3 is silenced by Sema3A-induced NSmiRNAs

A vastly accepted view posits that local translation in axons is triggered by stimuli, either by chemotropic and maturation cues during development or under injury conditions in adults (Batista and Hengst, 2016; Jung et al., 2012; Rangaraju et al., 2017). For instance, Sema3A induces the synthesis of proteins that elicit cytoskeletal remodelling and steering (Campbell and Holt, 2001; Wu et al., 2005). Here, however, we report the basal translation of APP, TUBB3 and *β*-Actin in individual axons elongating *ex vivo* on laminin substrate in absence of chemotropic and trophic cues. This is in agreement with several studies that have documented the basal translation of specific transcripts (Batista et al., 2017; Eng et al., 1999; Preitner et al., 2014; Taylor et al., 2013). A recent report furthermore revealed wide scale protein synthesis that occurs in isolated unstimulated *Xenopus laevis* axons within minutes (Cagnetta et al., 2018). These newly synthesized proteins represent one third of the total axonal proteome suggesting the existence of an unsuspected rich and complex basal translatome.

While the induction of global translation by chemotrophic cues is well established, very little is known about the fate of the basal translatome upon cue exposure. Here, using single axon FRAP of Venus translational reporter constructs, we reveal that Sema3A rapidly suppresses the basal translation of TUBB3 and APP *ex vivo*. This is in line with two other studies that have also measured cue-induced decreases in the translation of specific molecules in distal axons (Cagnetta et al., 2018; Yao et al., 2006). Overall, it is conceivable that two cue-activated pathways may co-exist in parallel to regulate the expression of two separate sets of proteins: a dominant pathway eliciting a burst of LPS and a secondary pathway inducing a trough of LPS or “LPS inhibition” (LPS-I). Both cue-induced LPS and LPS-I may ultimately lead to cytoskeleton remodelling and changes in growth cone behavior. LPS-I may be used as an alternative to proteasome degradation, which is not systematically employed for cue-mediated growth cone response (Campbell et al., 2001).

One key unresolved question is how cues inhibit basal LPS in axons. Here, we provide a series of evidence demonstrating that cue-induced LPS-I is mediated by NSmiRNAs. Using single axon FRAP of Venus-TUBB3-3’UTR constructs, we show that a cue-induced miRNA silences TUBB3 translation locally and that this effect is not due to a generic activation of miRNAs but to the cue-induced local biogenesis of miRNAs. Overall, our data suggest that an RNA-based signaling pathway exists, composed of mRNA and NSmiRNAs, two serially connected components of the same regulatory circuit. In response to cue, NSmiRNAs impinge on basal LPS of mRNAs to induce LPS-I. LPS and LPS-I are thereby coupled and coordinately regulated by NSmiRNAs acting on 3’UTR regulatory motifs. This coupling may generate a crucial leverage point for repellent cues to quickly and accurately adjust the desired level of individual proteins, including tubulin isotypes, within growth cones. Since Sema3A-induced growth cone turning does not depend on proteome degradation, contrary to other cues such as Netrin-1 (Campbell et al., 2001), this RNA-based mechanism may be crucial to regulate rapid changes in protein expression in response to specific repellents. To a large extent, this type of RNA-based signaling would allow to tightly control the rate and type of protein production for cytoskeletal remodelling, and thereby confer a higher order of regulatory potential to ensure the exquisite precision required for brain wiring.

What is the biological implication of the NSmiRNAs-triggered LPS-to-LPS-I switch for axon development? We uncover that this switch mediates cue-induced growth cone steering. We reveal that Sema3A triggers growth cone collapse response *ex vivo* and axon targeting *in vivo* through NSmiRNAs. NSmiRNAs, in turn, silence TUBB3 mRNA translation upon Sema3A exposure *ex vivo* and in the vicinity of Sema3A-expressing territories *in vivo*. Taken together, these data suggest that Sema3A-induced NSmiRNAs lead to the silencing of MT tubulin isotype TUBB3, MT depolymerization, and ultimately, growth cone steering. Collectively, our data thus support a model whereby Sema3A-produced NSmiRNAs elicit a rapid shift in axon behavior from axon elongation supported by basal TUBB3 LPS, to growth cone collapse prompted by TUBB3 LPS-I (Fig. S8F). According to this model, blocking NSmiRNA production prevents Sema3A-induced LPS-I of TUBB3 leading to the persistent production of TUBB3 and the maintenance of MT throughout the growth cone. As a consequence, growth cones fails to collapse *ex vivo* and, aberrantly continue to grow and meander *in vivo*, consistently with the phenotype that we observed.

In conclusion, our results reveal that inactive pre-miRNAs are actively transported to local sites for miRNA biogenesis and function, similarly to the subcellular translocation and subsequent local translation of silent mRNAs into functional proteins. At the growth cone, cue-induced NSmiRNAs impinge on local protein production by inhibiting basal LPS of their target mRNAs thereby contributing to changing growth cone direction. This type of ncRNA-based signaling pathway constitutes an additional regulatory layer that warrants the high degree of precision required for brain wiring. Many drug targets are detected within specific subcellular compartments. However, drug design does not often incorporate strategies for subcellular delivery (Rajendran et al., 2010). Since miRNA-based therapy using miRNA mimics and antimiRs are emerging as new promising therapeutics (Rupaimoole and Slack, 2017), our results lay important ground for the design of new clinical tools based on the targeted delivery and local activation of miRNAs.

## Contributions

**E.C.** carried out IF on Dicer transgenic mice, miRNA-seq analysis, axonal LCM, most RT-qPCR, some analyses of pre-miRNA and MB trafficking, MB and CD63 co-traffick experiments, collapse assay, all single axon FRAP experiments and analysis. **A.G.** performed MB design, pre-miRNA synthesis, most kymograph analysis of pre-miRNA and MB trafficking, thermal denaturation, microinjection, pre-miRNA KD, some RT-qPCR and MB quantification, pre-miRNA and MB co-localization experiments, Nocodazole treatment and live imaging. **S.St.** performed RNA-seq analysis and miRNA target prediction. **M.R.** performed MB design, pre-miRNA synthesis, and MSD analysis. **I.D.C.** carried out pre-miRNA and MB co-localization quantification, MB and CD63 co-trafficking experiments and analysis. **A.I.** performed stage 40 *in vivo* pathway analyses including rescue experiments with mimics. **S.B.** performed collapse assay, stage 44/45 *in vivo* pathway analysis and behavioral assay. **G.S.R.** performed stage 44/45 *in vivo* pathway analysis. **C.A.-G.** carried out miRNA-seq analysis. **E.C.** and **M.-L.B.** wrote the manuscript. All authors commented on the manuscripts. **E.C., M.R., C.A.-G., M.-L.B.** had supervisory roles. **M.-L.B.** supervised the project.

## Acknowledgements

The authors would like to thank Giovanna Berto for help with IF on transgenic mice, Benita Turner-Bridger for discussions on MB technology, Daniele Arosio, Angela Re and Silvia Pizzini for help with MSD analysis, Julie Lin and Hovy Ho-Wai Wong for help with *ex vivo* and *in vivo* single axon FRAP, C.E. Holt for granting access to HH lab Spinning disc confocal microscope, Institute for biomedicine EURAC imaging, CIBIO Imaging and MOF facilities. We also would like to thank Nick Ingoglia for stimulating discussions, Paolo Macchi and Giovanni Stefani for critically reading the ms. This study was supported by Erasmus+ Travel grant (E.C.), University of Trento PhD studentship (to A.I.), Marie Curie Career Integration (618969 GUIDANCE-miR), G. Armenise-Harvard Foundation Career, and MIUR SIR (RBSI144NZ4) grants (to M.-L.B.).

## ANIMAL MODEL MAINTENANCE

### *Xenopus laevis* embryos maintenance

*Xenopus laevis* embryos were obtained by *in vitro* fertilization, raised in 0.1x MMR diluted pH 7.5 at 14-22°C and staged according to Nieuwkoop and Faber (Nieuwkoop and Faber, 1994). All animal experiments were approved by the University of Trento Ethical Review Committee and by the Italian “Ministero della Salute” both according to the D.Lgs nr.116/92 and with the authorization n°1159/2016-PR and n°546/2017-PR according to art.31 of D.lgs. 26/2014.”

### Mus musculus

C57BL6 mice with a N-terminal Flag-HA2 tagged Dicer (Dcr^FH/FH^) were kindly donated by Dónal O’Carroll. Mice were housed and maintained, in accordance with the Decreto Legislativo 4 marzo 2014, n°26. All animal experiments were approved by the Italian “Ministero della Salute” with the authorization n°1159/2016-PR according to art.31 of D.lgs. 26/2014.”

## METHODS

### *In vitro* MB thermodynamics

To assess the thermodynamic characteristics of the MB, thermal denaturation profile of the MB was carried out, in which the fluorescence signal was recorded as a function of temperature, both in absence and presence of the target sequence. A thermocycler was adopted to carry out the thermal denaturation of the beacon, using “cold start” program: 15°C for 3 minutes, then from 16°C to 95°C increasing 1°C/minute.

### Immunohistochemistry on mice section and imaging

P0 brains and E13.5 whole heads were fixed in 4 % PFA (Life Technologies) overnight at 4°C, washed in 1xPBS (Gibco) and transferred into a 30 % sucrose (ACS reagent) solution in 1x PBS and left shaking at 4°C until sunk. Samples were embedded in OCT (Leica) and 14 µm sections were obtained with Leica CM 1850 UV cryostat.

Antigen retrieval was then performed in a preheated steamer for 25 minutes by immersion in pH6 10 mM sodium citrate (Sigma). Sections were permeabilized in TPBS (0.1 % Triton X-100 [Fisher Chemical] diluted in 1xPBS) and blocked with 10 % heat inactivated goat serum (NGS, Gibco) for 2 hours at room temperature. Rabbit anti-HA Y-11 (sc-805, 1:50, Santa Cruz Biotechnology) was used to stain Dicer-HA, mice anti-neurofilament (3A10, 1:500, Developmental Studies Hybridoma Bank, DSHB) was used to track neurons. Primary antibodies were incubated overnight at 4°C. Secondary antibodies (goat anti-rabbit Alexa Fluor 488 [A10684, 1:1000, Life technologies] and goat anti-mouse Alexa Fluor 594 [A11072, 1:1000, Life technologies]). Sections were counterstained with nuclear marker ToPro (T3605, 1:1000, Molecular probes) and mounted with ProLong Gold Antifade Mountant without DAPI (Molecular Probes).

Slides were acquired by confocal Leica TCS SP5 microscope or confocal Leica SP8 with white light lasers using a 40x objective. A zoom factor of 3 (Fig. 1D,S1C) or 1.99 (Fig. 1D) was applied during acquisition. All images were acquired through sequential scanning between frames. Gain and offset were optimized for best signal to noise ratio and the settings were maintained throughout acquisitions as well as the pinhole at 1.0 AU.

### *In situ* hybridization

DIG labeled LNA oligonucleotides complementary to *Xenopus laevis* miR-181a (5’-DigN -ACTCACCGACAGCGTTGAATGTT- DigN-3’), xtr-miR-181b (5’-DigN -CCCACCGACAGCAATGAATGTT- DigN-3’) and scrambled probes (5’- DigN -GTGTAACACGTCTATACGCCCA- DigN-3’) were obtained from Exiqon and added at 1 nM. 14 μm sections containing eye and brain of fixed stage 40 embryos were obtained using a Leica CM 1850 UV cryostat, collected on SuperFrost slides (Thermo Scientific) and processed for ISH as described previously (Baudet et al., 2012). Pre-hybridization step was performed at room temperature for 4 hours, followed by incubation of the hybridization solution containing denatured probe for 16 hours at 55°C. Slides were then imaged on upright widefield Zeiss Imager M2 fluorescence microscope equipped with AxioCam MRc color camera 5 Megapixel, using EC Plan Neofluar 20X/0.5 and EC Plan Neofluar 10X/0.3 objectives.

### Blastomere Microinjection

Morpholino mixtures were injected into both dorsal animal blastomeres of eight-cell-stage embryos as described previously (Baudet et al., 2012). The concentration and mixture used in each experiments are specified in the figure legends.

### Electroporation

RGC electroporation was performed as previously described (Baudet et al., 2012). After injection, the mixture were delivered at 18 V, by applying 8 electric pulses of 50 ms duration at 1000 ms intervals, except for in vivo FRAP experiment, where 2 pulses were applied instead of 8. The concentration and mixture used in each experiments are specified in the figure legends.

### Open book preparation

Open book preparation were obtained from brains as before (Baudet et al., 2012) acquired using a Leica DMi8 inverted fluorescence microscope. The z-stacks of serial images comprising the entire contralateral optic pathway were captured. Bright-field images were acquired as a reference of the tectum position.

### Behavioral assay

Embryos were grown at 18°C embryos until stage 44/45 on a white background in 12 h light/12 h dark environment. The behavioral assay was performed according to (Viczian and Zuber, 2014) with minor modifications. Embryos were grown into home made tanks whose walls were partitioned into black and white. A webcam (Logitech) was fixed to image the whole tank. One embryo was positioned at the time in the top black corner and the record started as soon as it was gently released. After 2 minutes the record was stopped, the orientation (180° turn) of the bottom black and white tank was inverted, and recording was restarted. The same embryo was sequentially tested 6 to 8 times.

### *Xenopus* retinal explants culture

Glass coverslips (Bellco) or glass-bottom dishes (MatTek) were subsequently coated with poly-L-lysine (Sigma, 10 μg/mL diluted in water) and with coated with laminin (Sigma, 10 μg/mL diluted in L-15 medium (Gibco). Eyes were dissected from anesthetized embryos, and cultured at 20°C for 20-24 hours in 60 % L-15 and 1 % Antibiotic-Antimycotic (ThermoFisher).

### Culture and *Xenopus* embryos fixation

Cultures were fixed in 2 % paraformaldehyde (PFA, Life Technologies), 7.5 % (wt/vol) sucrose (ACS reagent) for 30 minutes. Embryos were fixed in 4 % PFA (Life Technologies) for 2 hours at room temperature then rinsed three times in 1X PBS (Gibco).

### Collapse assay

200 ng/ml human recombinant Sema3A-FC (R&D System), 250 ng/ml Slit2 (R&D System) or PBS (for control) were bathed to explant culture for 10 minutes and then fixed. Collapse assay was performed as previously done (Bellon et al., 2017), and counted blind to the observer. For isolated axon preparations, axons closer than 100 μm to the cut were excluded from the count.

### Immunohistochemistry on *Xenopus laevis* and imaging

Fixed explants were permeabilized with 0.1 % Triton X-100 (Fisher Chemical). Blocking was carried out with 5 % heat inactivated goat serum (Gibco) diluted in TPBS for 1hr. Mouse monoclonal anti-Ago2 (a kind gift from Dónal O’Carroll, 1:50) and rabbit anti-Dicer (sc-30226, 1:50, Santa Cruz Biotechnology) were used. Primary antibodies were incubated overnight at 4°C.

Secondary antibodies Alexa 488 anti-mouse or 594 (F(ab’)2 fragments) anti-rabbit (Life Technologies) were used at 1:1000 and incubated 1 hour at room temperature in the dark. Explants were washed and mounted with Prolong Gold (Molecular Probes).

Zeiss observer Z1 inverted microscope equipped with AxioCam MR3, 1.4-megapixel monochromatic camera or an inverted Leica Dmi8 epifluorescence microscope equipped coupled with a sCMOS monochromatic camera (AndorZyla 4.2 Megapixel), were used with a magnification of HC Plan Apochromat CS2 63X/1.4 Oil objective.

### Laser capture microdissection (LCM) *ex vivo*

Stage 37/38 *Xenopus* embryos were cultured on RNase free POL (Polyester) membranes (Leica) with 1 mL 60 % L-15 medium (Gibco) for 24 hours. The following day, cultures were stained with FM-1-43FX dye (Thermo Fisher) for 20 minutes, fixed in 1 % PFA (Life Technologies) for 5 minutes and dehydrated in ethanol (Sigma) (25 %, 50 %, 75 %, 90 % and 100 %) for 1 minute each. Axons were captured using the Leica microdissector LMD6500 with the following settings: 20X and 40X (magnification), 33-38 (power), 1 (aperture), 16-14 (speed), 0 (specimen balance), 50 (offset).

### Isolated axon

Axons were severed from the explants at the stereomicroscope as before (Baudet et al., 2012). The complete experimental procedure was concluded within 1.5 hours after the first cut.

### Axonal transfection

NeuroMag Transfection Reagent (OZ Biosciences) were mixed and incubated for 20 minutes with MOs in a 1:100 proportion, added to the culture and incubated for 15 minutes on the magnetic base (OZ Biosciences) at RT. After transfection the plates were incubated for 30 minutes at 20°C, followed by three consecutive washes of 200 µL culture medium without perturbing the axons in the plate.

### RNA extraction

Single Cell RNA Purification Kit (Norgen) was used to extract total RNA from axonal samples following manufacturer’s instruction. In the lysis step, 200 µL of the lysis buffer (Buffer RL, Norgen) were mixed with *β*-mercaptoethanol (Sigma, 1:100), added to the axonal culture and incubated for 5 minutes. RNA was eluted by applying 9 µL Elution Solution twice to the column. Total RNA from eyes was extracted using Norgen Total RNA Purification Micro Kit (Norgen) following the manufacturer’s instructions and running an on-column RNAse free-DNAse I treatment (Norgen). For small fraction RNA collection the Split kit (Lexogen) was used, following manufacturer’s instruction. The integrity and concentration of RNA obtained from an isolated axon sample was assessed using the Agilent RNA 6000 Pico kit (Agilent technologies, Germany) on a Bioanalyzer (Agilent 2100 Bioanalyzer, Germany) for sensitive applications from small sample size (LCM & RNA-seq).

### RNA retrotranscription

3-5 µl axonal RNA from LCM or isolated axons, or 10 ng total RNA from eye samples were retrotranscribed with SuperScript^®^ IV (Thermofisher) using random hexamers primers (Euroclone) following the manufacturer’s instructions. Gene specific miRNA retrotranscription was performed with TaqMan™ MicroRNA Reverse Transcription Kit (Thermofisher) and the TaqMan qPCR assay (Thermofisher) following the manufacturer’s instructions.

### PCR and gel visualization

2 µl 1:3 diluted cDNA (from isolated axon preparation), or 3-4 µl undiluted cDNA (from LCM axonal collection) were used as input for GoTaq G2HotStart Polymerase (Promega) PCR reaction following manufacturer’s instruction. For mice genotyping, a small piece of tail was collected post-mortem, DNA was extracted by alkaline lysis and heating of the sample (Truett et al., 2000) and 4 µl of the supernatant were used directly as input for PCR reaction. Primers list and annealing temperature used are reported in the Reagent Table. PCR products were loaded on a 2 % TAE (Euroclone) or 3 % TBE (Thermofisher) agarose gel (Sigma Aldrich), run at 5.5 V/cm on an electrophoretic apparatus (BioRad) and visualized with UVITec Alliance LD2. The DNA was stained with Clear Sight DNA Stain (Bioatlas).

### RT-qPCR

Pre-miRNA expression level were investigated using Power SYBR Green PCR Master Mix (x2) (Applied Biosystem) and 0.25 µl of each 10 µM primer (all primers sequence are reported in Reagent Table), while TaqMan™ Universal Master Mix II (Thermofisher) was used for miRNAs amplification with the following TaqMan™ MicroRNA Assay (4427975 or 4440886): miR-181a-5p (ID: 000480); miR-181a-1-3p (ID: 004367_mat); miR-181a-2-3p (ID:005555_mat); miR-182 (ID: 000597) and snU6 (ID: 001973). qPCR cycling were run on BioRad CFX96 following manufacturer’s instructions.

### miR-181 target prediction and candidate selection

#### Identification of axonal RNA

10 ng of axonal RNA collected from isolated axons and SPRI beads (Beckman Coulter) purified were used as input for the Ovation SoLo RNA-Seq System, Custom AnyDeplete (NuGEN) and sequenced using NextSeq 500 - MID, paired-end 80nt approach (Illumina) at the EMBL Genomics Core facility (Heidelberg, Germany). Using the AnyDeplete system *Xenopus laevis* rRNA sequences identified through the ncbi nucleotide and the SILVA databases (Quast et al., 2012) were removed.

After ensuring raw sequence read quality with FastQC, reads were trimmed using Trimmomatic according to the Ovation SoLo RNA-Seq System instructions. Additionally, we removed adapter sequences and reads shorter than 40 bp. Trimmed reads were mapped with HISAT2 (Kim et al., 2015a), using default settings apart from –rna-strandness FR, to the *Xenopus* laevis genome (version 9.1) with an added rRNA contig and subsequently sorted using samtools (Li et al., 2009).

Using a custom *Xenopus laevis* transcriptome annotation, reads mapping to genes were quantified with FeatureCounts (Liao et al., 2014) using the following parameters: -p -t exon -g gene_id -s 1. The expression levels for each gene were calculated using the rpkm() function from the edgeR package (Robinson et al., 2010). We identified genes as axonally present if they were detectable at or above 1 FPKM.

#### miR-181 target prediction with TargetScan

*Xenopus laevis* miR-181a targets were predicted using custom scripts (TargetScan 6 (Garcia et al., 2011)) with custom annotated *Xenopus laevis* 3’UTR sequences. All predicted human and mouse miR-181a targets, including those with poorly conserved target sites but removing non-canonical ones, were downloaded from TargetScanHuman 7.1 and TargetScanMouse 7.1 (Garcia et al., 2011) respectively.

#### Identification of candidates

Axonally present predicted miR-181a-5p *Xenopus laevis* targets with an annotated 3’UTR of at least 50 bp length were screened using their human orthologs ncbi entrez IDs against the following Reactome (Fabregat et al., 2016) pathways of interest: axon guidance pathway (R-HSA-422475) and integrin cell surface interactions pathway (R-HSA-216083). In addition, we filtered the obtained predicted miR-181a-5p targets for those that are within the top 20 % of predicted targets and are also predicted to be targeted by miR-181-5p in human or mouse.

### Pre-miRNA candidate selection

Axonal pre-miRNAs candidates were selected from the published data [GEO accession number: <u>GSE86883</u>, (Bellon et al., 2017)]. The two libraries yielded 7.8 and 10.8 million reads. Candidate pre-miRNAs were identified by reads spanning the loop sequence of the pre-miRNA in addition to the -5p and -3p mature miRNA isoforms. The sequencing data were mapped to the available *Xenopus tropicalis, Xenopus laevis* and *Danio rerio* pre-miRNA sequences present in miRBase v21 (Bellon et al., 2017). For the pre-miRNAs candidate selection all sequences were required to perfectly match to *Xenopus laevis*. To achieve this aim all the reference sequences used in the mapping contained in miRBase v21 were blasted against the J-strain 9.2 Genome in Xenbase, using the default settings (E value 0.1, BLOSUM62 matrix), checking perfect match between reads and organism.

### Live imaging of exogenous and endogenous labeled pre-miR trafficking

Live imaging was performed inverted Leica Dmi8 epifluorescence microscope equipped coupled with a sCMOS monochromatic camera (AndorZyla 4.2 Megapixel) and with a HC PL Apo CS2 63x/1.4 immersion oil objective. The acquisition mode was set to 12-bit grayscale and “low noise” gain mode for fluorescence and “high well capacity” for phase contrast. No binning was applied to the acquisition. Exposure time and light intensity were chosen to optimize the signal to noise ratio, but were always kept invariant for the same batch of analysis. Exposure time was kept as low as possible (100-150 ms). Time-lapses for pre-miR-181a-1 trafficking analysis consisted of 150 to 300 consecutive frames recorded continuously, for a total time of 17 to 44 s, respectively with a 0.144 s delay between two consecutive frames, while it consisted of 130 frames per channel for co-trafficking with CD63. The distal end of single axons was chosen by the phase for imaging, strictly avoiding bundles, and had to comprise a stereotypical growth cone. This selected axon segment had to be at least 50-100 μm to the growth cone and at least 100 μm far from the soma.

### *Ex vivo* FRAP

For Fig. 8B-C,E-G and S8B, axons were imaged using a PerkinElmer Spinning Disk UltraVIEW ERS, Olympus IX81 inverted spinning disk confocal microscope, 60X UPLSAPO objectives (NA 1.3), equipped with Hamamatsu C11440-22CU camera. Axons were photobleached with a 488 nm laser with the following settings: “60” for PK cycles; “1” for PK step size; “8000ms” for PK spot period; “2” for PK spot cycles; “Small” for PK spot size; “None” for PK attenuation. For Fig. 8D, the labeled axons were visualized with an inverted TILL Photonics iMIC2, using a UPLSAPO 60x/1.2 water-immersion objective and a AVT Stingray F145B.30fps as detector. mRFP and Venus were visualized using an oligochrome Xenon arc lamp epifluorescence. Axons were photobleached with a 488 nm-laser using the following settings: dwell time (ms/µm^2^) “1”; scan line “optimum”; line overlapping “41 %”; ROI loop count “1”; experiment loop count “10”. Images for both mRFP and Venus were acquired before photobleaching, immediately after photobleaching, and then each minute for 10 minutes. Each image was captured with a Z-stack of 0.5 µm and an imaging depth of 2-3 µm. In control experiments, the protein synthesis was blocked with 100 µM cyclohexamide (CHX, Sigma) incubated for 30 minutes in the culture dishes before imaging. In the stimulated conditions with Sema3A, the cue was added to the cultures immediately after photobleaching.

### *In vivo* FRAP

Electroporated embryos with mRFP and Venus noUTR/Venus-TUBB3 3’UTR WT and MRE mutated were raised until stage 40/41 and prepared for live imaging as previously reported (Ströhl et al., 2017). The electroporated eye was removed in order to avoid trafficking from the soma to the RGC axonal compartment. Each image was captured with a Z-stack of 0.7 µm and an imaging depth of 7 µm.

## DNA AND OLIGOS CONSTRUCTS

### Plasmids

The plasmids pCS2-Venus, pCS2-Venus-ACTB-3’UTR and pCS2-mRFP were kind gifts from Christine Holt. The 3’UTRs of TUBB3 chL (Xelaev18022595m.g), APP chL (Xelaev18011533m.g), and THBS1 chL (Xelaev18042667m.g) were cloned into the pCS2 Venus plasmid between monomeric Venus coding sequence and polyA signal. RNA from stage 37/38 eyes was reverse transcribed with SuperScript III First-Strand Synthesis System (ThermoFisher) using 50 µM Oligo(dT) primers (Euroclone), and PCR reaction with Q5 High-Fidelity DNA Polymerase 2x Master mix (NEB) was performed following the manufacturer’s instructions. The mutated fragment after gel extraction and purification (QIAquick Gel Extraction Kit, Qiagen) were joint through overlapping-extension PCR (OE-PCR).

Two out of the three MREs in THBS1 3’ UTR were mutated through OE-PCR, while a third site, close to the 5’ part of the amplicon was inserted by Q5 mutagenesis kit (NEB) following manufacturer’s instructions. Primers used for wild type and mutated 3’ UTR amplification are reported in Table 5. Constructs thus obtained were sequenced. Mutated MREs of the same length as the WT and not matching any known axonal miRNA (Bellon et al., 2017) seed were selected.

Purified WT and mutated 3’UTR were ligated into the vector using T4 DNA Ligase Buffer (NEB) and high-efficiency NEB 5-alpha Competent E. coli cells (#C2987, NEB) were transformed following manufacturer’s instructions.

### *In vitro* synthesis and labeling of pre-miR-181a-1 transcript

The DNA template for the synthesis was obtained by elongation of two oligos (Table 5) complementary over 23 bp (100 µM each). The primers were annealed, elongated at 12°C using 25 U T4 DNA polymerase (NEB) and the obtained DNA fragment was purified with NucleoSpin PCR Clean-up kit (Macherey-Nagel).

The pre-miR *in vitro* synthesis was performed using T7 MEGAshortscriptTM kit (Ambion) from 1 µg purified DNA and following manufacturer’s instructions, including DNase TURBO digestion step to remove the template following transcription. To check pre-miR transcript integrity and length, 1 µl of the purified product was run on an 8M UREA (ACROS Organics) 10 % polyacrylamide (Euroclone) under denaturing conditions, using SYBR™ Gold Nucleic Acid Gel Stain 1000X (Life technologies).

5 µg of the *in vitro* synthesized pre-miR-181a-1 thus obtained was labeled with cy3 or cy5 using Label IT^®^ Nucleic Acid Labeling Kit (Mirus) following the manufacturer’s instructions.

### Molecular beacon

Labeled molecular beacon probes with sequence 5’- CA**UUG C{C}UUUA{G}AUAC CAA**UG -3’ was synthesized by Eurogentec (Belgium). The sequence in bold indicates the portion of the probe’s loop that is complementary to the pre-miR-181a-1 sequence. We used the *in silico* RNA and DNA folding prediction algorithm DinaMelt to design adequate thermodynamic characteristics (Markham and Zuker, 2005). We selected the GC content of the stem according to the recommended 30-60 % (Bratu et al., 2011), so that the melting temperature Tm of our MB was adapted to our working temperature (i.e. Tm 20-30°C higher than 20°C) (Simon et al., 2010; Søe et al., 2011), ensuring that, when unbound, it is in a closed conformation, stable and does not fluoresce whilst spontaneously anneals to its target. We designed a backbone composed of 2’-O-methyl ribose and included 2 centrally placed LNA nucleotides (bracket), as they improve avoiding nuclease digestion stability but also hybridization efficiency and probe specificity (Kierzek et al., 2005; Majlessi et al., 1998). MB was designed to maximize signal-to-noise ratio by choosing an appropriate fluorophore, cy3, and associated quencher “black hole quencher 2” (BHQ2) and a strong 5-nt stem with an adequate melting temperature (Tm) to ensure that the probe does not fluoresce when unbound or when used at 20°C, our working temperature (Baker et al., 2013; Tyagi and Alsmadi, 2004).

The reverse complement of the MB was blasted (www.xenbase.org) to check for possible off-targets in *Xenopus laevis*. The blast tool “blastn-DNA query to DNA database” was used, selecting as database “Xenopus Laevis J-strain 9.1 Genome” and as E value cut off 10. Blast against the whole genome was selected to investigate also possible off-targets on ncRNAs and the E value cut off was set at 10 to detect even the less probable off-targets. MB sequence or part of its sequence was complementary to additional 8 genomic loci but no corresponding reads were detected in axons by RNA-seq analysis. Therefore, MB does not match any known *Xenopus laevis* RNA sequence within the axon besides pre-miR-181a-1 (Table 2).

## QUANTIFICATION AND STATISTICAL ANALYSIS

### Statistic Analysis

All data were analyzed with Prism (GraphPad 6 or 7) and all experiments were performed in at least three independent biological replicates. For all tests, the significance level was α = 0.05. Exact number of replicates, tests used and statistics are reported in the Figure legend and in Table 3. All the statistics analysis pertaining to FRAP analysis are detailed in Table 4.

### Behavioral assay

Analysis of the movie acquired during the test were done measuring the amount of time the tadpole stays on the black side of the tank before reaching the white side. The time (in seconds) was expressed as percentage of the whole movie duration (at least 120 seconds) using *VLC software*.

### qPCR analysis

For RT-qPCR quantitative analysis, cycle threshold (Ct) were defined with CFX96 BioRad software v3.1, as mean of three technical replicates per sample. All technical replicates have a standard deviation smaller than 0.35. All Ct values are smaller than 35. Amplification efficiency of the new designed primers (pre-miR-181a-1, pre-miR-181a-2 and pre-miR-182) were investigated with standard curves independently from the actual experiments. To calculate miRNA or pre-miRNA differential expression, ΔCt method (Schmittgen et al., 2008) was applied as follows: 1/(2^[(CtmiR_X_-Ct_U6_)]).

### Life imaging analysis: kymograph

Movies were mapped back onto kymographs of the live pre-miR-181a-1, MB or CD63 movement using ImageJ. The image processing software FIJI, plugin “KymoReslicedWide” FIJI/ImageJ plugin was used to generate kymographs from timelapse movies, because of its high accuracy in detecting particle trajectories. For pre-miR-181a-1 trafficking studies, the specific macro tsp050706.txt for FIJI/imageJ software (author: J. Rietdorf FMI Basel + A. Seitz EMBL Heidelberg) was used, once the kymograph was obtained, to extrapolate from the tracked traces information about velocity and spatial directionality of the particles. The plugin was custom modified to also obtain particle directionality.

For co-trafficking analysis, the two kymographs (MB and CD63-eGFP puncta) were merged on and artificial color assigned using FIJI. Overlapping trace were considered as co-transported puncta when CD63 trace overlapped over the entire trajectory of the MB trace.

### MSD analysis

Region of interest (ROI) containing a segment from a single axon but not its growth cone was selected. Trajectories of single cy3-particles were recovered with Fiji/ImageJ using TrackMate plugin for automated single-particle tracking (Tinevez et al., 2017). To discriminate trajectories of particle that underwent directed motion from nonspecific noise and immobile objects, we selected trajectories based on their total displacement (minimum of 2.9 μm) and duration (80 consecutive frames). For each recovered trajectory, the mean square displacement (MSD) was calculated as follows: MSD (τ) = <(x(t+τ) – x(t))^2^ + (y(t+τ) – y(t))^2^> (Eq2) where x and y are the coordinates of the particle along the axon, t and τ are the absolute and lag times, respectively, and the brackets represent the time average. This calculation was performed for τ=25 % of the total time of the trajectory (Ruthardt et al., 2011).

The MSD data were fitted with an anomalous diffusion model: MSD = Aτ ^α^+B (Eq1), see Results section, where A depends on the motion properties of the particle, B is the residual MSD, and the coefficient α is an indication of the particle motion-type (Otero et al., 2014). Diffusion D was calculated according to D=MSD/qt (Eq3) where t is time and q is a constant depending on the dimension of the fitting model (q=2Xdim=4 in our case). Eq1 can be rewritten as t=r^2^/4D (Eq4) where t is time in second, and r is the displacement length in µm.

### FRAP

The mean intensity of the Venus signal normalized per growth cone area was measured with Volocity 64x software (Fig. 8B-C,E-G and S8B,) or ImageJ (Fig. 8D), by manually tracing the terminal of RGC axons on the sum of the z-stack for that specific timepoint. mRFP signal was used as reference to trace the axons. The background intensity was measured for the same area in proximity to the growth cone of interest and removed from the axonal Venus signal. From the background-corrected fluorescence intensity at each timepoint (F) the fluorescence signal after photobleaching (F_0_) was subtracted and normalized to the fluorescence signal pre-photobleaching: FRAP = (F-F_0_)/F_p_.

FRAP data were reported as curves of the recovered fluorescence signal in a 10-minute timeframe. The different curves were described by fitting the data with a nonlinear model (“one-phase decay” option in Prism) and differences between conditions were analyzed for statistical significance with an extra sum-of-square F test. Each axon acquired belonged to different eye explant. To avoid technical biases, the order of processing of the stimulation conditions with Sema3A, MO and co-MO samples, or WT and mutated were randomized both in term of organoculture and acquisition.

## KEY RESOURCES TABLE

- Table 1 - miR-181 targets identification in isolated axons
- Table 2 - MB Blast in Xenbase
- Table 3 - Stat table
- Table 4 - FRAP stat
- Table 5 - List of primers and oligos

## References

Alami, N.H., Smith, R.B., Carrasco, M.A., Williams, L.A., Winborn, C.S., Han, S.S.W., Kiskinis, E., Winborn, B., Freibaum, B.D., Kanagaraj, A., et al. (2014). Axonal Transport of TDP-43 mRNA Granules Is Impaired by ALS-Causing Mutations. Neuron 81, 536–543.

Aschrafi, A., Schwechter, A.D., Mameza, M.G., Natera-Naranjo, O., Gioio, A.E., and Kaplan, B.B. (2008). MicroRNA-338 Regulates Local Cytochrome c Oxidase IV mRNA Levels and Oxidative Phosphorylation in the Axons of Sympathetic Neurons. J. Neurosci. 28, 12581–12590.

Baker, M.B., Bao, G., and Searles, C.D. (2013). The use of molecular beacons to detect and quantify microRNA. Methods Mol. Biol. Clifton NJ 1039, 279–287.

Batista, A.F.R., and Hengst, U. (2016). Intra-axonal protein synthesis in development and beyond. Int. J. Dev. Neurosci. 55, 140–149.

Batista, A.F.R., Martínez, J.C., and Hengst, U. (2017). Intra-axonal Synthesis of SNAP25 Is Required for the Formation of Presynaptic Terminals. Cell Rep. 20, 3085–3098.

Baudet, M.-L., Zivraj, K.H., Abreu-Goodger, C., Muldal, A., Armisen, J., Blenkiron, C., Goldstein, L.D., Miska, E.A., and Holt, C.E. (2012). miR-124 acts through CoREST to control onset of Sema3A sensitivity in navigating retinal growth cones. Nat. Neurosci. 15, 29–38.

Baudet, M.-L., Bellon, A., and Holt, C.E. (2013). Role of microRNAs in Semaphorin function and neural circuit formation. Semin. Cell Dev. Biol. 24, 146–155.

Bauer, K.E., Kiebler, M.A., and Segura, I. (2017). Visualizing RNA granule transport and translation in living neurons. Methods 126, 177–185.

Baumann, S., Pohlmann, T., Jungbluth, M., Brachmann, A., and Feldbrügge, M. (2012). Kinesin-3 and dynein mediate microtubule-dependent co-transport of mRNPs and endosomes. J. Cell Sci. 125, 2740–52.

Bellon, A., Iyer, A., Bridi, S., Lee, F.C.Y., Ovando-Vázquez, C., Corradi, E., Longhi, S., Roccuzzo, M., Strohbuecker, S., Naik, S., et al. (2017). miR-182 Regulates Slit2-Mediated Axon Guidance by Modulating the Local Translation of a Specific mRNA. Cell Rep. 18, 1171–1186.

Bicker, S., Khudayberdiev, S., Weiss, K., Zocher, K., Baumeister, S., and Schratt, G. (2013). The DEAH-box helicase DHX36 mediates dendritic localization of the neuronal precursor-microRNA-134. Genes Dev. 27, 991–996.

Bielas, S.L., Serneo, F.F., Chechlacz, M., Deerinck, T.J., Perkins, G.A., Allen, P.B., Ellisman, M.H., and Gleeson, J.G. (2007). Spinophilin facilitates dephosphorylation of doublecortin by PP1 to mediate microtubule bundling at the axonal wrist. Cell 129, 579–91.

Bovolenta, P., and Mason, C. (1987). Growth cone morphology varies with position in the developing mouse visual pathway from retina to first targets. J. Neurosci. Off. J. Soc. Neurosci. 7, 1447–60.

Bratu, D.P., Catrina, I.E., and Marras, S.A.E. (2011). Tiny molecular beacons for in vivo mRNA detection. Methods Mol. Biol. Clifton NJ 714, 141–157.

Cabili, M.N., Dunagin, M.C., McClanahan, P.D., Biaesch, A., Padovan-Merhar, O., Regev, A., Rinn, J.L., and Raj, A. (2015). Localization and abundance analysis of human lncRNAs at single-cell and single-molecule resolution. Genome Biol. 16, 20.

Cagnetta, R., Frese, C.K., Shigeoka, T., Krijgsveld, J., and Holt, C.E. (2018). Rapid Cue-Specific Remodeling of the Nascent Axonal Proteome. Neuron 99, 29–46.e4.

Campbell, D.S., and Holt, C.E. (2001). Chemotropic responses of retinal growth cones mediated by rapid local protein synthesis and degradation. Neuron 32, 1013–1026.

Campbell, D.S., Regan, A.G., Lopez, J.S., Tannahill, D., Harris, W.A., and Holt, C.E. (2001). Semaphorin 3A elicits stage-dependent collapse, turning, and branching in Xenopus retinal growth cones. J. Neurosci. Off. J. Soc. Neurosci. 21, 8538–47.

Charrin, S., Jouannet, S., Boucheix, C., and Rubinstein, E. (2014). Tetraspanins at a glance. J. Cell Sci. 127, 3641–3648.

Chin, A., and Lécuyer, E. (2017). RNA localization: Making its way to the center stage. Biochim. Biophys. Acta Gen. Subj. 1861, 2956–2970.

Davis, L., Dou, P., DeWit, M., and Kater, S.B. (1992). Protein synthesis within neuronal growth cones. J. Neurosci. Off. J. Soc. Neurosci. 12, 4867–77.

Dent, E.W., and Gertler, F.B. (2003). Cytoskeletal dynamics and transport in growth cone motility and axon guidance. Neuron 40, 209–27.

Eng, H., Lund, K., and Campenot, R.B. (1999). Synthesis of beta-tubulin, actin, and other proteins in axons of sympathetic neurons in compartmented cultures. J. Neurosci. Off. J. Soc. Neurosci. 19, 1–9.

Fabregat, A., Sidiropoulos, K., Garapati, P., Gillespie, M., Hausmann, K., Haw, R., Jassal, B., Jupe, S., Korninger, F., McKay, S., et al. (2016). The Reactome pathway Knowledgebase. Nucleic Acids Res. 44, D481–D487.

Falk, J., Konopacki, F.A., Zivraj, K.H., and Holt, C.E. (2014). Rab5 and Rab4 Regulate Axon Elongation in the Xenopus Visual System. J. Neurosci. 34, 373–391.

Földes-Papp, Z., König, K., Studier, H., Bückle, R., Breunig, H.G., Uchugonova, A., and Kostner, G.M. (2009). Trafficking of mature miRNA-122 into the nucleus of live liver cells. Curr. Pharm. Biotechnol. 10, 569–78.

Garcia, D.M., Baek, D., Shin, C., Bell, G.W., Grimson, A., and Bartel, D.P. (2011). Weak seed-pairing stability and high target-site abundance decrease the proficiency of lsy-6 and other microRNAs. Nat. Struct. Mol. Biol. 18, 1139–1146.

Gershoni-Emek, N., Altman, T., Ionescu, A., Costa, C.J., Gradus-Pery, T., Willis, D.E., and Perlson, E. (2018). Localization of RNAi Machinery to Axonal Branch Points and Growth Cones Is Facilitated by Mitochondria and Is Disrupted in ALS. Front. Mol. Neurosci. 11, 311.

Gibbings, D.J., Ciaudo, C., Erhardt, M., and Voinnet, O. (2009). Multivesicular bodies associate with components of miRNA effector complexes and modulate miRNA activity. Nat. Cell Biol. 11, 1143–1149.

Godement, P., Salaün, J., and Imbert, M. (1984). Prenatal and postnatal development of retinogeniculate and retinocollicular projections in the mouse. J. Comp. Neurol. 230, 552–575.

Grishok, A., Pasquinelli, A.E., Conte, D., Li, N., Parrish, S., Ha, I., Baillie, D.L., Fire, A., Ruvkun, G., and Mello, C.C. (2001). Genes and mechanisms related to RNA interference regulate expression of the small temporal RNAs that control C. elegans developmental timing. Cell 106, 23–34.

Hancock, M.L., Preitner, N., Quan, J., and Flanagan, J.G. (2014). MicroRNA-132 Is Enriched in Developing Axons, Locally Regulates Rasa1 mRNA, and Promotes Axon Extension. J. Neurosci. 34, 66–78.

Hengst, U., Cox, L.J., Macosko, E.Z., and Jaffrey, S.R. (2006). Functional and Selective RNA Interference in Developing Axons and Growth Cones. J. Neurosci. 26, 5727–5732.

Holt, C.E., and Harris, W.A. (1983). Order in the initial retinotectal map in Xenopus: a new technique for labelling growing nerve fibres. Nature 301, 150–2.

Hutvagner, G., McLachlan, J., Pasquinelli, A.E., Bálint, E., Tuschl, T., and Zamore, P.D. (2001). A Cellular Function for the RNA-Interference Enzyme Dicer in the Maturation of the let-7 Small Temporal RNA. Science 293, 834–838.

Iyer, A.N., Bellon, A., and Baudet, M.-L. (2014). microRNAs in axon guidance. Front. Cell. Neurosci. 8, 78.

Jung, H., Yoon, B.C., and Holt, C.E. (2012). Axonal mRNA localization and local protein synthesis in nervous system assembly, maintenance and repair. Nat. Rev. Neurosci. 13, 308–324.

Kar, A.N., MacGibeny, M.A., Gervasi, N.M., Gioio, A.E., and Kaplan, B.B. (2013). Intra-axonal Synthesis of Eukaryotic Translation Initiation Factors Regulates Local Protein Synthesis and Axon Growth in Rat Sympathetic Neurons. J. Neurosci. 33, 7165–7174.

Kierzek, E., Ciesielska, A., Pasternak, K., Mathews, D.H., Turner, D.H., and Kierzek, R. (2005). The influence of locked nucleic acid residues on the thermodynamic properties of 2′-O-methyl RNA/RNA heteroduplexes. Nucleic Acids Res. 33, 5082–5093.

Kim, E., and Jung, H. (2015). Local protein synthesis in neuronal axons: why and how we study. BMB Rep. 48, 139–146.

Kim, D., Langmead, B., and Salzberg, S.L. (2015a). HISAT: a fast spliced aligner with low memory requirements. Nat. Methods 12, 357–360.

Kim, H.H., Kim, P., Phay, M., and Yoo, S. (2015b). Identification of precursor microRNAs within distal axons of sensory neuron. J. Neurochem. 134, 193–199.

Konopacki, F.A., Wong, H.H.-W., Dwivedy, A., Bellon, A., Blower, M.D., and Holt, C.E. (2016). ESCRT-II controls retinal axon growth by regulating DCC receptor levels and local protein synthesis. Open Biol. 6, 150218.

Kos, A., Loohuis, N.O., Meinhardt, J., Bokhoven, H. van, Kaplan, B.B., Martens, G.J., and Aschrafi, A. (2016). MicroRNA-181 promotes synaptogenesis and attenuates axonal outgrowth in cortical neurons. Cell. Mol. Life Sci. 73, 3555–3567.

Kye, M.-J., Liu, T., Levy, S.F., Xu, N.L., Groves, B.B., Bonneau, R., Lao, K., and Kosik, K.S. (2007). Somatodendritic microRNAs identified by laser capture and multiplex RT-PCR. RNA 13, 1224–1234.

Lee, Y.S., Pressman, S., Andress, A.P., Kim, K., White, J.L., Cassidy, J.J., Li, X., Lubell, K., Lim, D.H., Cho, I.S., et al. (2009). Silencing by small RNAs is linked to endosomal trafficking. Nat. Cell Biol. 11, 1150–1156.

Leung, K.-M., Lu, B., Wong, H.H.-W., Lin, J.Q., Turner-Bridger, B., and Holt, C.E. (2018). Cue-Polarized Transport of β-actin mRNA Depends on 3′UTR and Microtubules in Live Growth Cones. Front. Cell. Neurosci. 12, 300.

Li, H., Handsaker, B., Wysoker, A., Fennell, T., Ruan, J., Homer, N., Marth, G., Abecasis, G., Durbin, R., and 1000 Genome Project Data Processing Subgroup (2009). The Sequence Alignment/Map format and SAMtools. Bioinformatics 25, 2078–2079.

Liao, Y., Smyth, G.K., and Shi, W. (2014). featureCounts: an efficient general purpose program for assigning sequence reads to genomic features. Bioinformatics 30, 923–930.

Liu, X.-A., Rizzo, V., and Puthanveettil, S.V. (2012). PATHOLOGIES OF AXONAL TRANSPORT IN NEURODEGENERATIVE DISEASES. Transl. Neurosci. 3, 355–372.

Lugli, G., Torvik, V.I., Larson, J., and Smalheiser, N.R. (2008). Expression of microRNAs and their precursors in synaptic fractions of adult mouse forebrain. J. Neurochem. 106, 650–661.

Maday, S., Twelvetrees, A.E., Moughamian, A.J., and Holzbaur, E.L.F. (2014). Axonal Transport: Cargo-Specific Mechanisms of Motility and Regulation. Neuron 84, 292–309.

Magdesian, M.H., Gralle, M., Guerreiro, L.H., Beltrão, P.J.I., Carvalho, M.M.V.F., Santos, L.E. da S., de Mello, F.G., Reis, R.A.M., and Ferreira, S.T. (2011). Secreted human amyloid precursor protein binds semaphorin 3a and prevents semaphorin-induced growth cone collapse. PloS One 6, e22857.

Majlessi, M., Nelson, N.C., and Becker, M.M. (1998). Advantages of 2’-O-methyl oligoribonucleotide probes for detecting RNA targets. Nucleic Acids Res. 26, 2224–9.

Markham, N.R., and Zuker, M. (2005). DINAMelt web server for nucleic acid melting prediction. Nucleic Acids Res. 33, W577–W581.

Much, C., Auchynnikava, T., Pavlinic, D., Buness, A., Rappsilber, J., Benes, V., Allshire, R., and O’Carroll, D. (2016). Endogenous Mouse Dicer Is an Exclusively Cytoplasmic Protein. PLOS Genet. 12, e1006095.

Natera-Naranjo, O., Aschrafi, A., Gioio, A.E., and Kaplan, B.B. (2010). Identification and quantitative analyses of microRNAs located in the distal axons of sympathetic neurons. RNA 16, 1516–1529.

Nieuwkoop, P.D. (Pieter D.., and Faber, J. (1994). Normal table of Xenopus laevis (Daudin) : a systematical and chronological survey of the development from the fertilized egg till the end of metamorphosis (New York : Garland Pub).

Otero, M.G., Alloatti, M., Cromberg, L.E., Almenar-Queralt, A., Encalada, S.E., Pozo Devoto, V.M., Bruno, L., Goldstein, L.S.B., and Falzone, T.L. (2014). Fast axonal transport of the proteasome complex depends on membrane interaction and molecular motor function. J. Cell Sci. 127, 1537–1549.

Poirier, K., Saillour, Y., Bahi-Buisson, N., Jaglin, X.H., Fallet-Bianco, C., Nabbout, R., Castelnau-Ptakhine, L., Roubertie, A., Attie-Bitach, T., Desguerre, I., et al. (2010). Mutations in the neuronal \s s-tubulin subunit TUBB3 result in malformation of cortical development and neuronal migration defects. Hum. Mol. Genet. 19, 4462–73.

Pols, M.S., and Klumperman, J. (2009). Trafficking and function of the tetraspanin CD63. Exp. Cell Res. 315, 1584–92.

Preitner, N., Quan, J., Nowakowski, D.W., Hancock, M.L., Shi, J., Tcherkezian, J., Young-Pearse, T.L., and Flanagan, J.G. (2014). APC Is an RNA-Binding Protein, and Its Interactome Provides a Link to Neural Development and Microtubule Assembly. Cell 158, 368–382.

Quast, C., Pruesse, E., Yilmaz, P., Gerken, J., Schweer, T., Yarza, P., Peplies, J., and Glöckner, F.O. (2012). The SILVA ribosomal RNA gene database project: improved data processing and web-based tools. Nucleic Acids Res. 41, D590–D596.

Rajendran, L., Knölker, H.-J., and Simons, K. (2010). Subcellular targeting strategies for drug design and delivery. Nat. Rev. Drug Discov. 9, 29–42.

Rangaraju, V., tom Dieck, S., and Schuman, E.M. (2017). Local translation in neuronal compartments: how local is local? EMBO Rep. 18, 693–711.

Resovi, A., Pinessi, D., Chiorino, G., and Taraboletti, G. (2014). Current understanding of the thrombospondin-1 interactome. Matrix Biol. 37, 83–91.

Robinson, M.D., McCarthy, D.J., and Smyth, G.K. (2010). edgeR: a Bioconductor package for differential expression analysis of digital gene expression data. Bioinformatics 26, 139–140.

Roher, A.E., Kokjohn, T.A., Clarke, S.G., Sierks, M.R., Maarouf, C.L., Serrano, G.E., Sabbagh, M.S., and Beach, T.G. (2017). APP/Aβ structural diversity and Alzheimer’s disease pathogenesis. Neurochem. Int. 110, 1–13.

Rupaimoole, R., and Slack, F.J. (2017). MicroRNA therapeutics: towards a new era for the management of cancer and other diseases. Nat. Rev. Drug Discov. 16, 203–222.

Ruthardt, N., Lamb, D.C., and Bräuchle, C. (2011). Single-particle Tracking as a Quantitative Microscopy-based Approach to Unravel Cell Entry Mechanisms of Viruses and Pharmaceutical Nanoparticles. Mol. Ther. 19, 1199–1211.

Salogiannis, J., and Reck-Peterson, S.L. (2017). Hitchhiking: A Non-Canonical Mode of Microtubule-Based Transport. Trends Cell Biol. 27, 141–150.

Sambandan, S., Akbalik, G., Kochen, L., Rinne, J., Kahlstatt, J., Glock, C., Tushev, G., Alvarez-Castelao, B., Heckel, A., and Schuman, E.M. (2017). Activity-dependent spatially localized miRNA maturation in neuronal dendrites. Science 355, 634–637.

Santangelo, P., Nitin, N., LaConte, L., Woolums, A., and Bao, G. (2006). Live-cell characterization and analysis of a clinical isolate of bovine respiratory syncytial virus, using molecular beacons. J. Virol. 80, 682–688.

Schmittgen, T.D., Lee, E.J., Jiang, J., Sarkar, A., Yang, L., Elton, T.S., and Chen, C. (2008). Real-time PCR quantification of precursor and mature microRNA. Methods San Diego Calif 44, 31–38.

Simon, B., Sandhu, M., and Myhr, K.L. (2010). Live FISH: Imaging mRNA in living neurons. J. Neurosci. Res. 88, 55–63.

Smalheiser, N.R. (2008). Synaptic enrichment of microRNAs in adult mouse forebrain is related to structural features of their precursors. Biol. Direct 3, 44.

Søe, M.J., Møller, T., Dufva, M., and Holmstrøm, K. (2011). A Sensitive Alternative for MicroRNA In Situ Hybridizations Using Probes of 2′-O-Methyl RNA + LNA. J. Histochem. Cytochem. 59, 661–672.

Ströhl, F., Lin, J.Q., Laine, R.F., Wong, H.H.-W., Urbančič, V., Cagnetta, R., Holt, C.E., and Kaminski, C.F. (2017). Single Molecule Translation Imaging Visualizes the Dynamics of Local β-Actin Synthesis in Retinal Axons. Sci. Rep. 7, 709.

Taylor, A.M., Wu, J., Tai, H.-C., and Schuman, E.M. (2013). Axonal translation of β-catenin regulates synaptic vesicle dynamics. J. Neurosci. Off. J. Soc. Neurosci. 33, 5584–9.

Tinevez, J.-Y., Perry, N., Schindelin, J., Hoopes, G.M., Reynolds, G.D., Laplantine, E., Bednarek, S.Y., Shorte, S.L., and Eliceiri, K.W. (2017). TrackMate: An open and extensible platform for single-particle tracking. Methods 115, 80–90.

Tischfield, M.A., Baris, H.N., Wu, C., Rudolph, G., Van Maldergem, L., He, W., Chan, W.-M., Andrews, C., Demer, J.L., Robertson, R.L., et al. (2010). Human TUBB3 Mutations Perturb Microtubule Dynamics, Kinesin Interactions, and Axon Guidance. Cell 140, 74–87.

Truett, G.E., Heeger, P., Mynatt, R.L., Truett, A.A., Walker, J.A., and Warman, M.L. (2000). Preparation of PCR-quality mouse genomic DNA with hot sodium hydroxide and tris (HotSHOT). BioTechniques 29, 52, 54.

Turner-Bridger, B., Jakobs, M., Muresan, L., Wong, H.H.-W., Franze, K., Harris, W.A., and Holt, C.E. (2018). Single-molecule analysis of endogenous β-actin mRNA trafficking reveals a mechanism for compartmentalized mRNA localization in axons. Proc. Natl. Acad. Sci. U. S. A. 115, E9697–E9706.

Tyagi, S., and Alsmadi, O. (2004). Imaging native beta-actin mRNA in motile fibroblasts. Biophys. J. 87, 4153–62.

Vargas, J.N.S., Kar, A.N., Kowalak, J.A., Gale, J.R., Aschrafi, A., Chen, C.-Y., Gioio, A.E., and Kaplan, B.B. (2016). Axonal localization and mitochondrial association of precursor microRNA 338. Cell. Mol. Life Sci. 73, 4327–4340.

Viczian, A.S., and Zuber, M.E. (2014). A Simple Behavioral Assay for Testing Visual Function in *Xenopus laevis*. J. Vis. Exp.

Wang, B., and Bao, L. (2017). Axonal microRNAs: localization, function and regulatory mechanism during axon development. J. Mol. Cell Biol. 9, 82–90.

Wang, B., Pan, L., Wei, M., Wang, Q., Liu, W.-W., Wang, N., Jiang, X.-Y., Zhang, X., and Bao, L. (2015). FMRP-Mediated Axonal Delivery of miR-181d Regulates Axon Elongation by Locally Targeting Map1b and Calm1. Cell Rep. 13, 2794–807.

Wong, H.H.-W., Lin, J.Q., Ströhl, F., Roque, C.G., Cioni, J.-M., Cagnetta, R., Turner-Bridger, B., Laine, R.F., Harris, W.A., Kaminski, C.F., et al. (2017). RNA Docking and Local Translation Regulate Site-Specific Axon Remodeling In Vivo. Neuron 95, 852–868.e8.

Wu, K.Y., Hengst, U., Cox, L.J., Macosko, E.Z., Jeromin, A., Urquhart, E.R., and Jaffrey, S.R. (2005). Local translation of RhoA regulates growth cone collapse. Nature 436, 1020–1024.

Yamada, M., and Sekiguchi, K. (2015). Molecular Basis of Laminin–Integrin Interactions. In Current Topics in Membranes, pp. 197–229.

Yamaguchi, Y., and Miura, M. (2015). Programmed Cell Death in Neurodevelopment. Dev. Cell 32, 478–490.

Yao, J., Sasaki, Y., Wen, Z., Bassell, G.J., and Zheng, J.Q. (2006). An essential role for beta-actin mRNA localization and translation in Ca2+-dependent growth cone guidance. Nat. Neurosci. 9, 1265–73.

You, X., Vlatkovic, I., Babic, A., Will, T., Epstein, I., Tushev, G., Akbalik, G., Wang, M., Glock, C., Quedenau, C., et al. (2015). Neural circular RNAs are derived from synaptic genes and regulated by development and plasticity. Nat. Neurosci. 18, 603–610.

Zhang, Y., Ueno, Y., Liu, X.S., Buller, B., Wang, X., Chopp, M., and Zhang, Z.G. (2013). The MicroRNA-17-92 cluster enhances axonal outgrowth in embryonic cortical neurons. J. Neurosci. Off. J. Soc. Neurosci. 33, 6885–94.

Zivraj, K.H., Tung, Y.C.L., Piper, M., Gumy, L., Fawcett, J.W., Yeo, G.S.H., and Holt, C.E. (2010). Subcellular Profiling Reveals Distinct and Developmentally Regulated Repertoire of Growth Cone mRNAs. J. Neurosci. 30, 15464–15478.

